# Essential omega-3 fatty acids tune microglial phagocytosis of synaptic elements in the developing brain

**DOI:** 10.1101/744136

**Authors:** C. Madore, Q. Leyrolle, L. Morel, J.C. Delpech, A.D. Greenhalgh, C. Lacabanne, C. Bosch-Bouju, J. Bourel, A. Thomazeau, K.E. Hopperton, S. Beccari, A. Sere, A. Aubert, V. De Smedt-Peyrusse, C. Lecours, K. Bisht, L. Fourgeaud, S. Gregoire, L. Bretillon, N. J. Grant, J. Badaut, P. Gressens, A. Sierra, O. Butovsky, M.E. Tremblay, R.P. Bazinet, C. Joffre, A. Nadjar, S. Layé

**Affiliations:** Univ. Bordeaux, INRA, Bordeaux INP, NutriNeuro, UMR 1286, F-33000, Bordeaux, France; Ann Romney Center for Neurologic Diseases, Department of Neurology, Brigham and Women’s Hospital, Harvard Medical School, Boston, MA, USA; NeuroDiderot, Inserm, Université de Paris Diderot, F-75019 Paris, France; Department of Nutritional Sciences, University of Toronto, Toronto, M5S 3E2, Canada; Achucarro Basque Center for Neuroscience, University of the Basque Country and Ikerbasque Foundation, Leioa, 48940, Spain; Neurosciences Axis, CRCHU de Québec-Université Laval, Québec City, Québec, Canada; Molecular Neurobiology Laboratory, The Salk Institute for Biological Studies, La Jolla, California 92037, USA; Centre des Sciences du Goût et de l’Alimentation, AgroSup Dijon, CNRS, INRA, Univ. Bourgogne Franche-Comté, F-21000 Dijon, France; CNRS UPR3212, Institut des Neurosciences Cellulaires et Intégratives, Strasbourg, France; CNRS UMR5287, University of Bordeaux, Bordeaux, France; Centre for the Developing Brain, Department of Division of Imaging Sciences and Biomedical Engineering, King’s College London, King’s Health Partners, St. Thomas’ Hospital, London, SE1 7EH, United Kingdom; Evergrande Center for Immunologic Diseases, Brigham and Women’s Hospital, Harvard Medical School, Boston, MA, USA

**Author notes:** Co-first authors (C. Madore, Q. Leyrolle, L. Morel). Co-senior authors (A. Nadjar, S. Layé).

## Abstract

Omega-3 fatty acids (n-3 polyunsaturated fatty acids; n-3 PUFAs) are essential for the functional maturation of the brain. Westernization of dietary habits in both developed and developing countries is accompanied by a progressive reduction in dietary intake of n-3 PUFAs. Low maternal intake of n-3 PUFAs has been linked to neurodevelopmental diseases in epidemiological studies, but the mechanisms by which a n-3 PUFA dietary imbalance affects CNS development are poorly understood. Active microglial engulfment of synaptic elements is an important process for normal brain development and altered synapse refinement is a hallmark of several neurodevelopmental disorders. Here, we identify a molecular mechanism for detrimental effects of low maternal n-3 PUFA intake on hippocampal development. Our results show that maternal dietary n-3 PUFA deficiency increases microglial phagocytosis of synaptic elements in the developing hippocampus, through the activation of 12/15- lipoxygenase (LOX)/12-HETE signaling, which alters neuronal morphology and affects cognition in the postnatal offspring. While women of child bearing age are at higher risk of dietary n-3 PUFA deficiency, these findings provide new insights into the mechanisms linking maternal nutrition to neurodevelopmental disorders.

**One Sentence Summary:** Low maternal omega-3 fatty acids intake impairs microglia-mediated synaptic refinement *via* 12-HETE pathway in the developing brain.

## INTRODUCTION

Lipids are one of the main constituents of the central nervous system (CNS). Among these, arachidonic acid (AA, 20:4n-6) and docosahexaenoic acid (DHA, 22:6n-3) are the principal forms of the long chain (LC) polyunsaturated fatty acids, omega-6 and omega-3 (referred to as n-6 or n-3 PUFAs) of the grey matter (Sastry, 1985). These two LC PUFAs are found predominantly in the form of phospholipids and constitute the building blocks of brain cell membranes^1, 2^. AA and DHA are either biosynthesized from their respective dietary precursors, linoleic acid (LA, 18:2n-6) and α-linolenic acid (ALA, 18:3n-3), or directly sourced from the diet (mainly meat and dairy products for AA, fatty fish for DHA)^3, 4^. In addition, AA and DHA are crucial for cell signaling through a variety of bioactive mediators. These mediators include oxylipins derived from cyclooxygenases (COX), lipoxygenases (LOX) and cytochrome P450 (CYP) pathways^1^ that are involved in the regulation of inflammation and phagocytic activity of immune cells^5–11^. An overall increase in the dietary LA/ALA ratio and a reduction in LC n-3 PUFA intake, as found in the Western diet, leads to reduced DHA and increased AA levels in the brain^4, 12^.

DHA is essential for the functional maturation of brain structures^13^. Low n-3 PUFA consumption globally^14^ has raised concerns about its potential detrimental effects on the neurodevelopment of human infants^15^ and the incidence of neurodevelopmental diseases such as Autism Spectrum Disorder (ASD) and schizophrenia^16, 17^.

To understand the mechanisms by which n-3/n-6 PUFA imbalance affects CNS development, we investigated the impact of maternal dietary n-3 PUFA deficiency on offspring’s microglia, the resident immune cells involved in CNS development and homeostasis^18^. Microglia are efficient phagocytes of synaptic material and apoptotic cells, which are key processes in the developing brain^19–23^. Specifically, once neuronal circuits are established, microglia contribute to the refinement of synaptic connections by engulfing synaptic elements in early post-natal life, in a fractalkine- and complement cascade-dependent manner^24–29^. This refinement is important for behavioral adaptation to the environment^27, 30–33^.

We have previously reported that a maternal dietary n-3 PUFA deficiency alters the microglial phenotype and reduces their motility in the developing hippocampus^34^. Therefore, we investigated the effect of maternal (i.e. gestation and lactation) n-3 PUFA deficiency on microglial function and its consequences on hippocampal development. Using multidisciplinary approaches, we reveal that maternal exposure to an n-3 PUFA deficient diet impairs offspring’s microglial homeostatic signature and phagocytic activity, and neuronal circuit formation, resulting in behavioral abnormalities. N-3 PUFA deficiency drives excessive microglial phagocytosis of synaptic elements through the oxylipin 12/15-LOX/12-HETE pathway and is a key mechanism contributing to microglia-mediated synaptic remodeling in the developing brain. These results highlight the impact of bioactive lipid mediators on brain development and identify imbalanced nutrition as a potent environmental risk factor for neurodevelopmental disorders.

## RESULTS

### Maternal dietary n-3 PUFA deficiency alters the morphology of hippocampal neurons and hippocampal-mediated spatial working memory

The hippocampus plays an essential role in learning and memory, with the neurons of the CA1 region required for spatial working memory^35^. Therefore, we assessed the effects of maternal dietary n-3 PUFA deficiency on neuronal morphology in the offspring hippocampus CA1 using Golgi-Cox staining at post-natal day (P)21. Dendritic spine density of Golgi-stained CA1 pyramidal neurons was significantly decreased in n-3 deficient mice compared to n-3 sufficient mice (Figure 1A-B). This was associated with a decrease in the length of dendrites, whereas the complexity of dendritic arborization was unchanged (Figure 1A-B). Western blot analyses on whole hippocampi showed a significant decrease in the expression of the post-synaptic scaffold proteins PSD-95 and Cofilin, but not of SAP102, in n-3 deficient *vs.* n-3 sufficient mice (Figure 1C-D). AMPA and NMDA subunit expression levels were unaffected (Supplementary Figure 1A-B). In the Y-maze task, a hippocampus-dependent test assessing spatial working memory, n-3 deficient mice were unable to discriminate between the novel and familiar arms of the maze, unlike the n-3 sufficient mice (Figure 1E). These data show that maternal n-3 PUFA deficiency disrupts early-life (P21) spine density, neuronal morphology and alters spatial working memory.

**Figure 1:**
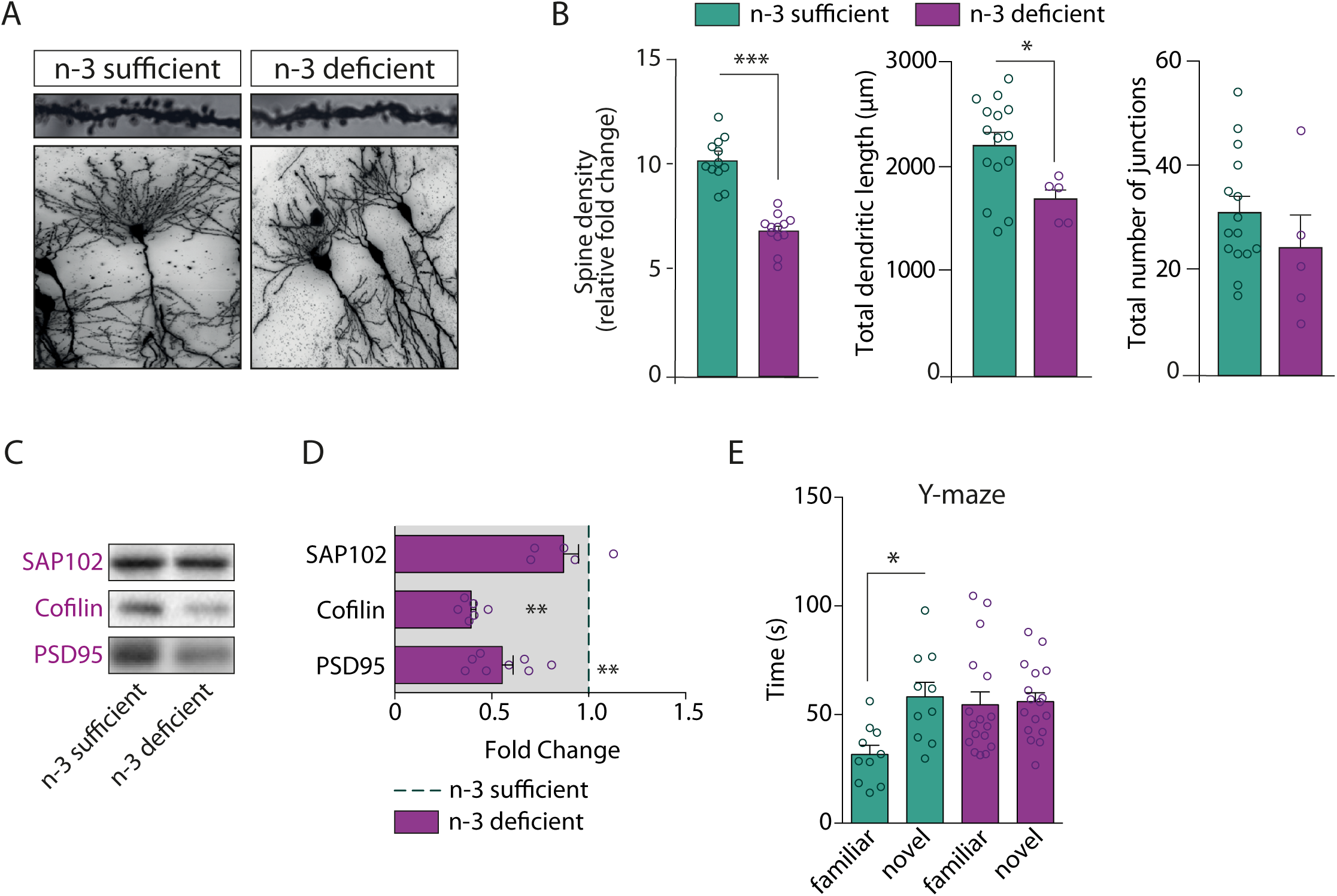
Maternal n-3 PUFA deficiency alters neuronal morphology and function. **A.** Representative images of tertiary apical dendrites (upper panel) and dendritic arborization (lower panel) of hippocampal CA1 pyramidal neurons from n-3 sufficient and n-3 deficient mice at P21 (Golgi staining). **B.** Quantification of spine density, total dendritic length and total number of junctions in n-3 deficient and n-3 sufficient animals. For spine counting, n=12 CA1 pyramidal neurons of the hippocampus, 15 to 45 segments per animal, 3 mice per group. Means ± SEM. Two-tailed unpaired Student’s t-test, t=6.396, ***p<0.0001. For dendritic arborization and total dendritic length, n=5 neurons from 3 mice for n-3 deficient mice and n=15 neurons from 4 mice for n-3 sufficient mice. Two-tailed unpaired Student’s t-test; Dendritic length, t=2.501, *p=0.0223; number of junctions: t=1.118, p=0.28. **C.** Representative Western blots. **D.** Expression of scaffolding proteins in n-3 deficient mice relative to n-3 sufficient mice. Means ± SEM; n=5-8 mice per group. Two-tailed unpaired Student’s t-test, t=0.9986, p= 0.35, SAP102; t=3.572, **p=0.0044, cofilin; t=3.529, **p=0.0042, PSD95. **E.** Time spent in novel *vs* familiar arm in the Y maze task. Means ± SEM; n=10-17 mice per group. Two-way ANOVA followed by Bonferroni *post-hoc* test: diet effect, F(1,50)=5.47, p=0.071; arm effect, F(1,50)=10.08, p=0.015; interaction, F(1,50)=8.08, p=0.029; familiar vs novel arm for n-3 diet mice: *p<0.05.

### Maternal dietary n-3 PUFA deficiency increases microglia-mediated synaptic loss

As microglia phagocytose pre- and post-synaptic elements to shape neural networks^24, 25, 27–29^, we used electron microscopy (EM) to analyze, at high spatial resolution, microglia–synapse interactions in the hippocampal CA1 region of n-3 deficient *vs.* n-3 sufficient mice (Figure 2A-B). At P21, *i.e*. at the peak of hippocampal synaptic pruning^36^, we found significantly more contacts between Iba1-positive microglial processes and the synaptic cleft and more dendritic spine inclusions within those processes in n-3 deficient mice (Figure 2B). Microglial processes from n-3 deficient mice contained more inclusions within endosomes such as spines and other cellular elements, along with a reduced accumulation of debris in the extracellular space, thus suggesting an increase in phagocytic activity^27^ (Figure 2A-B). This was not due to changes in microglial process size, as diet did not affect process morphology (Supplementary Figure 2A). In addition, confocal imaging and 3D reconstruction of microglia showed that PSD95-immunoreactive puncta within Iba1-positive microglia were more abundant in the hippocampus of n-3 deficient mice, corroborating the EM data (Figure 2C-D). To provide further support to this increase in microglial phagocytosis, we stimulated *ex vivo* hippocampal microglia from n-3 deficient *vs.* n-3 sufficient P21 mice with pHrodo E.Coli bioparticles. The results revealed that the percentage of phagocytic microglia is significantly higher in n-3 deficient mice across the time points studied (Figure 2E-F; Supplementary Figure 2B). Increased microglial phagocytosis in the hippocampus was not a compensation to a decrease in microglial density, as there were no differences in microglial density assessed by stereological counting in the different hippocampal sub-regions, including the CA1 (Supplementary Figure 2C-D).

**Figure 2:**
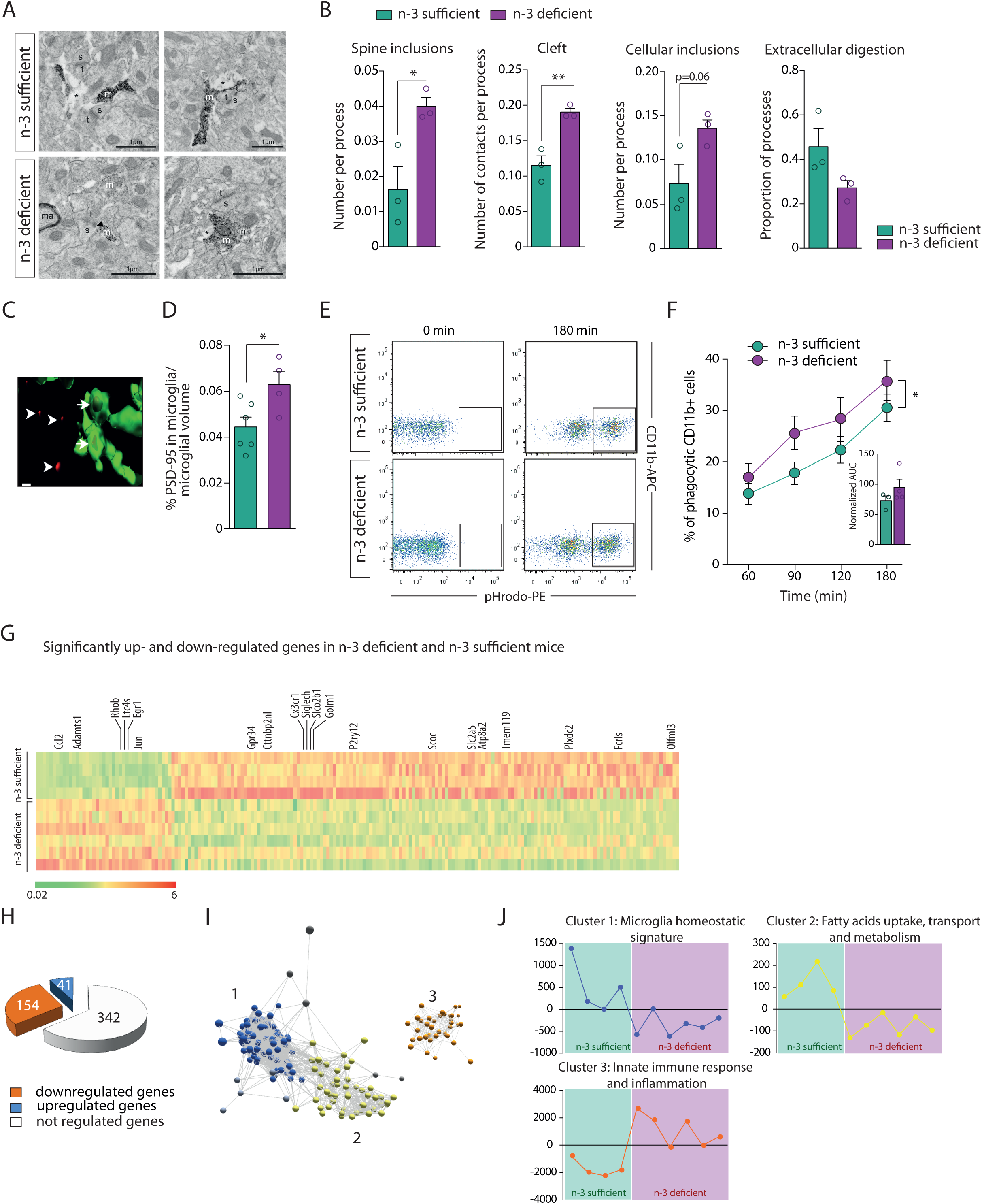
Maternal n-3 PUFA deficiency increases microglial phagocytosis and gene expression profile. **A-B.** Representative images (**A**) and quantification (**B**) of EM data from the CA1 region of n-3 sufficient and n-3 deficient mice. ‘s’= spine, ‘m’= microglial processes, ‘t’= terminals, *= extracellular debris, ‘ma’=myelinated axons. Arrowheads point to synaptic clefts. Scale bars=c1 μm. Two-tailed unpaired Student’s t-test; t=5.24, **p= 0.0063, cleft; t=3.366, *p=0.0282, spine inclusions; t=2.601, p=0.06, cellular inclusions; t=2.119, p=0.1015, extracellular digestion. **C.** Three-dimensional reconstructions of PSD95-immunoreactivity (red) outside (arrowheads) or colocalized with (arrows) small-diameter microglial processes tips (green). **D.** Quantification of PSD95 and Iba1 immunoreactivity colocalization in the CA1 region of n-3 deficient and n-3 sufficient group. Means ± SEM; n=4-6 mice per group. Two-tailed unpaired Student’s t-test; t=2.583, *p=0.032. **E.** Representative bivariate dot plots of isolated microglial cells gated on CD11b^+^/CD45^low^ expression from n-3 sufficient and n-3 deficient mice. **F.** FACS analysis of phagocytic uptake of pHrodo-labeled E. Coli bioparticles by CD11b^+^ microglial cells from both dietary groups after 180 min of incubation (expressed as the percent of cells that are CD11b^+^ and PE positive). Means ± SEM; n=3-4 per condition. Two-way ANOVA; diet effect: F(1, 20)=5.628, *p=0.028; time effect, F(3,20)=10.04, p=0.0003; Interaction, F(3,20)=0.1771, p=0.91. The area under the curve is presented as an independent graph in C. Two-tailed unpaired Student’s t-test, t=1.289, p=0.25. **G.** Heatmap of significantly up or downregulated genes in each dietary group. Each lane represents one animal (n=4 or 6 per group). The 20 homeostatic microglia unique genes that were significantly affected by the diet are labelled. **H.** Number of microglial genes that are up-regulated, down-regulated or not regulated by the diet. **I.** A transcript-to-transcript correlation network graph of transcripts significantly differentially expressed by diet groups was generated in Miru (Pearson correlation threshold *r* ≥ 0.85). Nodes represent transcripts (probe sets), and edges represent the degree of correlation in expression between them. The network graph was clustered using a Markov clustering algorithm, and transcripts were assigned a color according to cluster membership. **J.** Mean expression profile of all transcripts within clusters 1, 2 and 3.

Enhanced microglial phagocytic activity was also not linked to an increase in cell death, as the volume of the hippocampus, number of neurons and astrocytes, number of apoptotic cells and expression of pro-apoptotic protein Bax and anti-apoptotic protein Bcl-2 were not different between diets (Supplementary Figure 3A-F). Furthermore, it is unlikely that n-3 PUFA deficiency-induced increases in phagocytosis resulted from brain infiltration of peripherally-derived macrophages^37^. This is consistent with the observation that maternal n-3 PUFA deficiency did not modulate the number of CD11b-high/CD45-high cells within the whole brain (Supplementary Figure 3G) and that the blood-brain barrier was intact in both diet groups as revealed by the expression of Claudin-5 and GFAP as well as IgG extravasation (Supplementary Figure 3H-M).

### Maternal dietary n-3 PUFA deficiency dysregulates microglial homeostasis

During brain development, microglia display a unique transcriptomic signature that supports their physiological functions, including in synaptic refinement^38^. Therefore, we investigated whether and how n-3 PUFA deficiency modulates the homeostatic molecular signature of microglia isolated from n-3 deficient *vs*. n-3 sufficient mouse brains. Using a Nanostring-based mRNA chip containing 550 microglia-enriched genes^39^, 154 of these genes were found to be significantly down-regulated and 41 genes up-regulated in microglia from n-3 deficient mice whole brains (Figure 2G-J, Table 3). Among the 40 genes reported to be unique to homeostatic microglia by ^39^, 20 were significantly dysregulated, indicating that maternal n-3 PUFA deficiency alters homeostatic functions of microglia (Figure 2G). We next sought to define the diet-specific microglial phenotype by assessing patterns of gene co-expression using unbiased network analysis software, Miru^40^. The utility of identifying gene expression patterns and transcriptional networks underpinning common functional pathways in microglia has been previously described^41^. Using a Markov clustering algorithm to non-subjectively subdivide our data into discrete sets of co-expressed genes, we found three major clusters of genes that were differentially regulated in n-3 deficient mice (Figure 2I-J). The mean expression profiles of these three clusters showed that clusters 1 (microglial homeostatic signature) and 2 (fatty acid uptake, transport and metabolism) contained genes whose expression was relatively lower in the n-3 deficient group as compared to n-3 sufficient group. In contrast, cluster 3 (innate immune response and inflammation) contained genes with relatively greater expression in n-3 deficient mice (Figure 2J). These results show that an n-3 PUFA deficient diet dysregulates microglial homeostasis and increases microglial immune and inflammatory pathways during development.

### Lipid changes in microglia, not in synaptic elements, exacerbate phagocytosis after maternal dietary n-3 PUFA deficiency

N-3 PUFA deficient diets cause lipid alterations in whole brain tissue^42^, yet it is unclear to what extent individual cell types or cell compartments incorporate these changes. To assess this, we profiled the lipid composition of microglia and synaptosomes isolated from n-3 sufficient and n-3 deficient mice (Figure 3A-B). We observed that maternal n-3 PUFA deficiency significantly altered the lipid profile of both microglia and synaptosomes, with an increase in n-6 fatty acids and a decrease in n-3 fatty acids (Figure 3A-B). Moreover, DHA was decreased in microglia from n-3 deficient mice (Figure 3A), while C22:5n-6 (or DPA n-6), the n-6-derived structural equivalent of DHA, was increased as a marker of n-3 PUFA deficiency^12, 43^ (Figure 3A). Total AA was not significantly modified by the diet in microglia (Figure 3A), unlike total AA levels in the whole hippocampus (Supplementary Figure 4). In synaptosomes, total AA level was not changed but total DHA level was significantly decreased and DPA n-6 inversely increased, as observed in microglia (Figure 3B). Synaptosomes contain a wider array of fatty acids and more of them were significantly modified by the diet, such as the saturated fatty acid (SFA) stearic acid (18:0), and the monounsaturated fatty acid (MUFA) oleic acid (18:1 n-9). Overall, our results show that lipid profiles of both microglia and synaptosomes are altered by low maternal n-3 PUFA intake, displaying similarities and discrepancies (Figure 3A-B).

**Figure 3:**
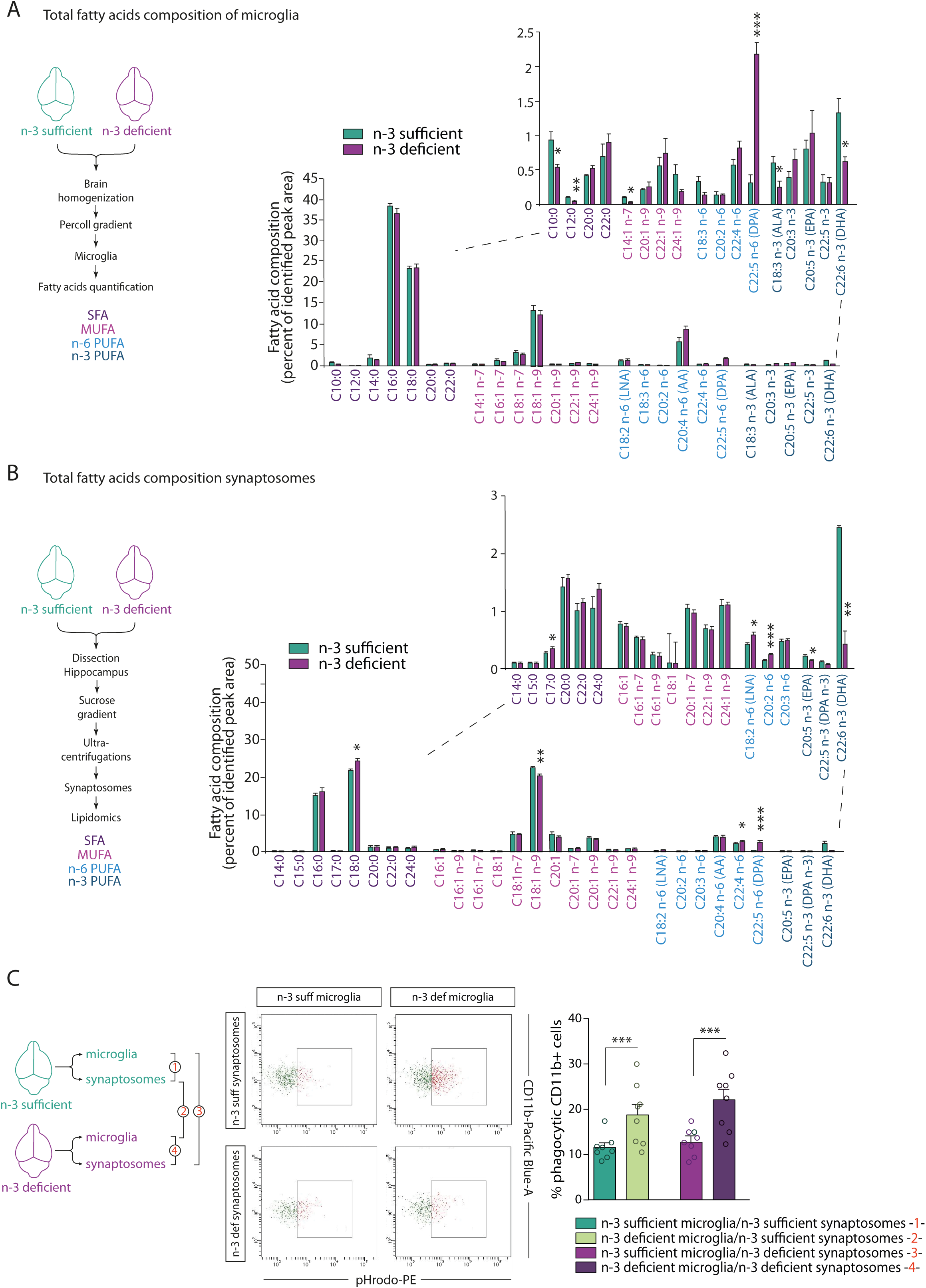
Maternal dietary n-3 PUFA deficiency exacerbates microglial phagocytosis of synaptic elements by predominantly impacting microglia. **A**. Fatty acid composition of microglial cells sorted from n-3 sufficient and n-3 deficient mice. Means ± SEM; n=4 mice per group. Insert: higher magnification of low expressed fatty acids. Two-tailed unpaired Student’s t-test; t=3.37, *p=0.015, C10:0; t=4.852, **p=0.0028, C12:0; t=2.473, *p=0.0482, C14:1 n-7; t=9.646, ***p<0.0001, DPA n-6; t=2.558, *p=0.043, ALA; t=2.972, *p=0.0249, DHA. **B.** Fatty acid composition of synaptosomes sorted from n-3 sufficient and n-3 deficient mice. Means ± SEM; n=4 mice per group. Insert: higher magnification of low expressed fatty acids. Two-tailed unpaired Student’s t-test; t=2.82, *p=0.047, C17:0; t=3.05, *p=0.038, C18:0; t=5.05, **p=0.0072 C18:1 n-9; t=2.99, *p=0.04, C18:2 n-6; t=26, ***p<0.0001, C20:2 n-6; t=3.57, *p=0.023, C22:4 n-6; t=12.68, ***p=0.0002, DPA n-6; t=4.49, *p=0.011, C20:5 n-3; t=7.12, **p=0.0021, DHA**. C.** FACS analysis of phagocytic uptake of pHrodo-labeled synaptosomes (sorted from n-3 sufficient or n-3 deficient mice) by CD11b^+^ microglial cells (sorted from n-3 sufficient or n-3 deficient mice). On FACS plots: x-axis= (pHrodo intensity (PE intensity), y-axis=SSC. Means ± SEM; n=8 per condition. Two-way ANOVA: microglial lipid status effect, F(1,28)=19.77, ***p=0.0001; synaptosomes lipid status effect, F(1,28)=1.63, p=0.2123; interaction, F(1,28)=0.2826, p=0.599.

To directly evaluate the effect of microglial and synaptosome lipid alterations on microglial phagocytic activity, we developed an *ex vivo* assay in which we exposed freshly isolated n-3 sufficient or n-3 deficient microglia to n-3 sufficient or n-3 deficient pHrodo-labelled synaptosomes (Figure 3C). We measured microglial phagocytic activity based on the level of cellular accumulated pHrodo fluorescence (Figure 3C). Phagocytosis was significantly increased in microglia from n-3 deficient mice, independent of the lipid profile of the synaptosome (Figure 3C). These data show that both microglial and synaptosome lipid profiles are modified by maternal n-3 PUFA deficiency, but changes in microglial lipid composition drive the exacerbation of microglia-mediated synaptic loss.

### Enhancement of microglia-mediated synaptic refinement by maternal PUFA deficiency is phosphatidyl serine (PS) recognition-independent

Microglia possess a number of phagocytic receptors to eliminate neuronal elements, notably by binding to externalized phosphatidyl serine (PS) at the surface of neurons^19, 44, 45^. Since PUFAs are the main constituents of membrane phospholipids, we assessed whether microglial phagocytic activity was enhanced through phosphatidyl serine (PS) recognition (Figure 4). First, we showed that cell-surface PS, stained with annexin-V, was unchanged between diets in the hippocampus (Figure 4A). We also quantified the expression levels of the microglial PS-binding protein milk fat globule EGF factor 8 (MFG-E8), which binds and activates a vitronectin receptor (VNR), TAM receptor tyrosine kinases (MER and Axl), triggering receptor expressed on myeloid cells-2 (TREM2) and CD33^19, 44, 45^. Only MFG-E8 expression level was significantly increased by maternal n-3 PUFA dietary deficiency (Figure 4B-G). We administered a blocking peptide cRGD which inhibits the interaction between MFG-E8 and VNR *vs* a scrambled control (sc-cRGD), to standard chow-fed animals and quantified the expression of the post-synaptic protein PSD-95. Under standard conditions, inhibiting MFG-E8 activity increased PSD-95 protein expression (Supplementary Figure 5). However, when the same approach was used in n-3 deficient *vs* n-3 sufficient mice, blocking MFG-E8 action on microglia did not prevent the reduction in PSD-95 expression in the hippocampus of n-3 deficient mice (Figure 4H). These results indicate that modulation of the microglial phagocytic capacity by early-life PUFAs is PS recognition-independent.

**Figure 4:**
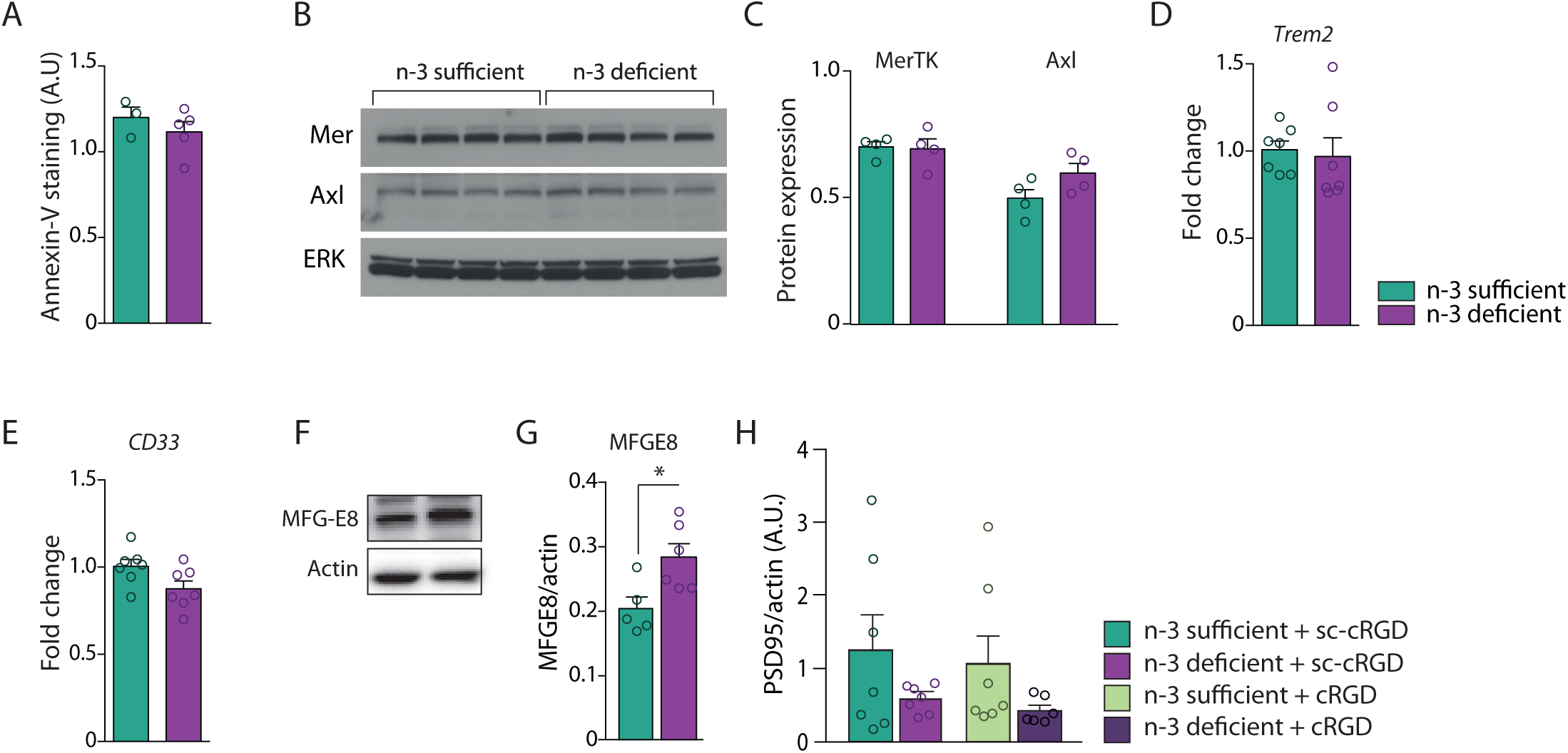
Maternal n-3 PUFA deficiency enhances microglial phagocytic capacity in a PS recognition-independent manner. **A.** Annexin V staining is not different between n-3 deficient and n-3 sufficient mice. Means ± SEM; n=3-5 mice per group. Two-tailed unpaired Student’s t-test; t=0.911, p= 0.397. **B.** Representative Western blots for Mer and Axl proteins. **C.** Quantification of Mer and Axl protein in −3 deficient and n-3 sufficient mice. Means ± SEM; n=4 mice per group. Two-tailed unpaired Student’s t-test; t=0.169, p=0.87, Mer; t=1.824, p=0.12, Axl. **D-E.** RT-qPCR detects the relative abundance of *trem2* (**D**) and *cd33* (**E**) mRNA in hippocampus of n-3 deficient and n-3 sufficient mice. Means ± SEM; n=7 mice per group. Two-tailed unpaired Student’s t-test; t=0.329, p=0.75, *trem2*; t=2.158, p=0.052, *cd33*. **F.** Representative Western blots for MFG-E8 protein. **G.** Quantification of MFG-E8 protein level in n-3 deficient vs n-3 sufficient mice. Means ± SEM; n=5-6 mice. Two-tailed unpaired Student’s t-test; t=2.803, *p=0.0206. **H.** Quantification of PSD95 protein expression in n-3 sufficient *vs* n-3 deficient mice treated with cRGD or its control (scrambled peptide sc-cRGD). Two-way ANOVA: diet effect, F(1,23)=4.017, p=0.057; treatment effect, F(1,23)=0.3417, p=0.56; interaction, F(1,23)=1.24×10^−5^, p=0.99.

### Maternal n-3 PUFA deficiency alters neuronal morphology and working spatial memory in a complement-dependent manner

Recently, the classical complement cascade has been described as a key mediator of microglia synaptic refinement in the developing brain^25, 46, 47^. Moreover, our EM data revealed more spine inclusions within microglia and more contacts between these cells and the synaptic cleft (Figure 2A-B). Thus, we explored the implication of the complement cascade in the exacerbation of microglia-mediated synaptic refinement observed in n-3 deficient mice. Protein expression levels of CD11b (a subunit of CR3) and C1q were significantly increased in the hippocampus CA1 and DG in n-3 deficient mice (Figure 5A-B, D-E). CD11b gene expression was also enhanced in these mice (Supplementary Figure 6C). Cleavage of C3 not only releases the opsonin C3b, but can also release the anaphylactic peptide C3a, which modulates microglial phagocytosis by acting on the receptor C3aR^48^. Our analyses also revealed a significantly increased density of C3aR immunostaining in the whole hippocampus of n-3 deficient mice, corroborating microglia transcriptomic data (Table 3). (Figure 5C,F; Table 3). The C3aR-positive cells were uniformly distributed throughout the hippocampus, including the CA1. Expression levels of *Cx3cr1*, *Cx3cl1* and *Tgfb*, all involved in microglia-mediated synaptic pruning^24, 25, 49^, were found to be increased in the whole hippocampus of n-3 deficient mice (Supplementary Figure 6). Finally, we quantified the C3 protein on freshly extracted hippocampus synaptosomes from n-3 deficient and n-3 sufficient mice using an ELISA assay. We found higher amounts of C3 under n-3 PUFA deficiency (Figure 5G). These findings indicate that low n-3 PUFA intake alters the developmental expression of complement cascade proteins both in microglia and at the synapse.

**Figure 5.**
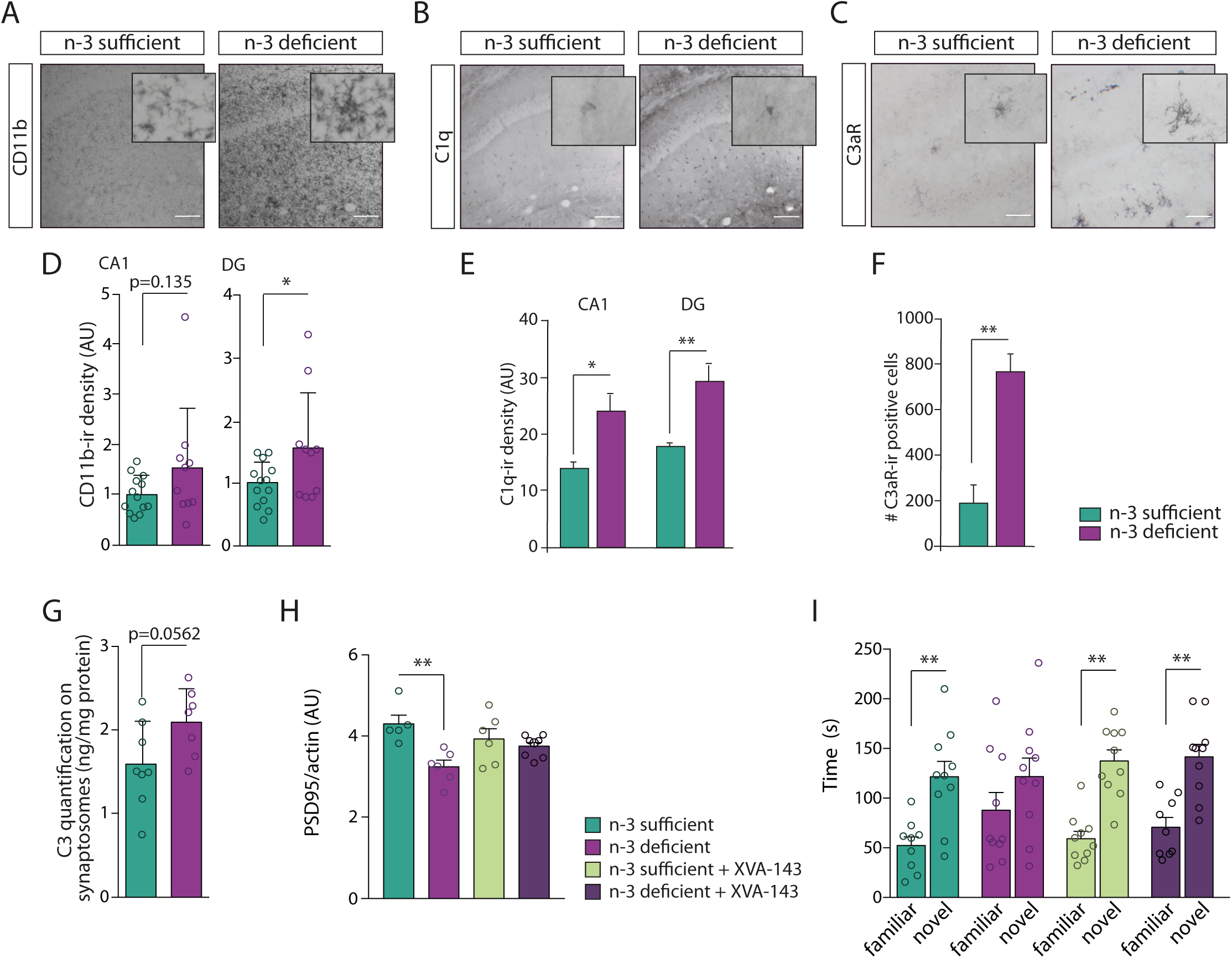
Maternal n-3 PUFA deficiency exacerbates microglia-mediated shaping of neuronal networks in a complement-dependent manner. **A-C.** Immunostaining for CD11b, C1q and C3aR in coronal sections of n-3 deficient and n-3 sufficient mice hippocampus. Scale bar=100μm. **D-F.** Quantification of CD11b, C1q and C3aR immunoreactivity in the hippocampus (CA1 and DG regions) of n-3 deficient and n-3 sufficient mice. Means ± SEM; n=3-13 mice per group. Two-tailed unpaired Student’s t-test; t=1.553, p=0.135, CD11b CA1; t=2.114, *p=0.466, CD11b DG; t=0.5495, *p=0.0223 C1q CA1; t=3.938, **p=0.0056, C1q DG; t=4.91, **p=0.0027, C3aR, total hippocampus. **G.** Protein quantification (ELISA) reveals that synaptosomes from n-3 deficient mice express more C3 than n-3 sufficient animals. Means ± SEM; n=7-8 mice per group; Two-tailed unpaired Student’s t-test; t=2.097, p=0.0562. **H.** Quantification of PSD95 protein expression in n-3 sufficient vs n-3 deficient mice treated with XVA-143 or its vehicle. Means ± SEM; n=5-8. Two-way ANOVA followed by Bonferroni *post-hoc* test: diet effect, F(1,21)=11.89, p=0.0024; treatment effect, F(1,21)=0.1436, p=0.708; interaction, F(1,21)=5.977, p=0.0234; n-3 sufficient vs n-3 deficient, **p<0.01. **I.** Time spent in novel *vs* familiar arm in the Y maze task in n-3 sufficient vs n-3 deficient mice treated with XVA-143 or its vehicle. Means ± SEM; n=9-10 mice per group. Paired t-test: n-3 sufficient group, **p=0.0019; n-3 deficient group, p=0.357; n-3 sufficient + XVA-143 group, **p=0.0013; n-3 deficient + XVA-143 group, **p=0.0099.

To test whether proper neuronal function could be restored by acting on the complement pathway *in vivo*, we antagonized CR3 activation in n-3 deficient mice. We first confirmed that a CR3 antagonist XVA-143, blocked n-6 PUFA (AA) induced increases in phagocytosis *in vitro* (Supplementary Figure 7A-D). We then administered the CR3 antagonist, XVA-143^50^ *in vivo* in the hippocampus four days before assessing the expression of the post-synaptic protein PSD95 at P21. Antagonizing CR3 prevented PSD-95 expression decrease in the hippocampus of n-3 deficient mice (Figure 5H). Injection of XVA-143 also restored optimal memory abilities in n-3 deficient mice in the Y-maze task (Figure 5I). These results show that complement activation increases microglia-mediated synaptic refinement when dietary n-3 PUFAs are low. Thus, a global increase in C3/CR3 interaction may account for the altered neuronal function observed in the hippocampus of n-3 deficient mice.

### Maternal n-3 PUFA deficiency alters microglial PUFA metabolism in the offspring

Our data suggest that n-3 PUFA deficiency specifically drives exacerbated microglial phagocytosis (Figure 3). Therefore, we assessed whether microglial PUFA metabolism was driving their phagocytic activity. Microglia isolated from n-3 deficient mice had higher levels of free, unesterified, AA and lower quantities of free EPA, as determined by high-performance liquid chromatography with tandem mass spectrometry (LC-MS-MS) (Figure 6A). When unesterified from the cellular membrane, free PUFAs are rapidly converted to bioactive mediators (or oxylipins) with potential effects on microglial cells^1, 7, 8, 51–54^. Interestingly, microglia from n-3 deficient animals presented greater levels of AA-derived bioactive mediators and lower amounts of DHA- and EPA-derived mediators (Figure 6B). Four AA-derived oxylipins were significantly increased by maternal n-3 PUFA deficiency: trioxilins TRXA3 and TRXB3, 8-HETE, 12-HETE, while three EPA-derived species were significantly decreased: +/−11-HEPE, 15S-HEPE and LTB5 and DHA-derived 17SHDoHE was also reduced (p=0.0957) (Figure 6C). Hence, we show for the first time the post-natal microglia oxylipin signature in the context of low maternal n-3 PUFA intake and that it is modified by the dietary content in n-3 PUFAs.

**Figure 6:**
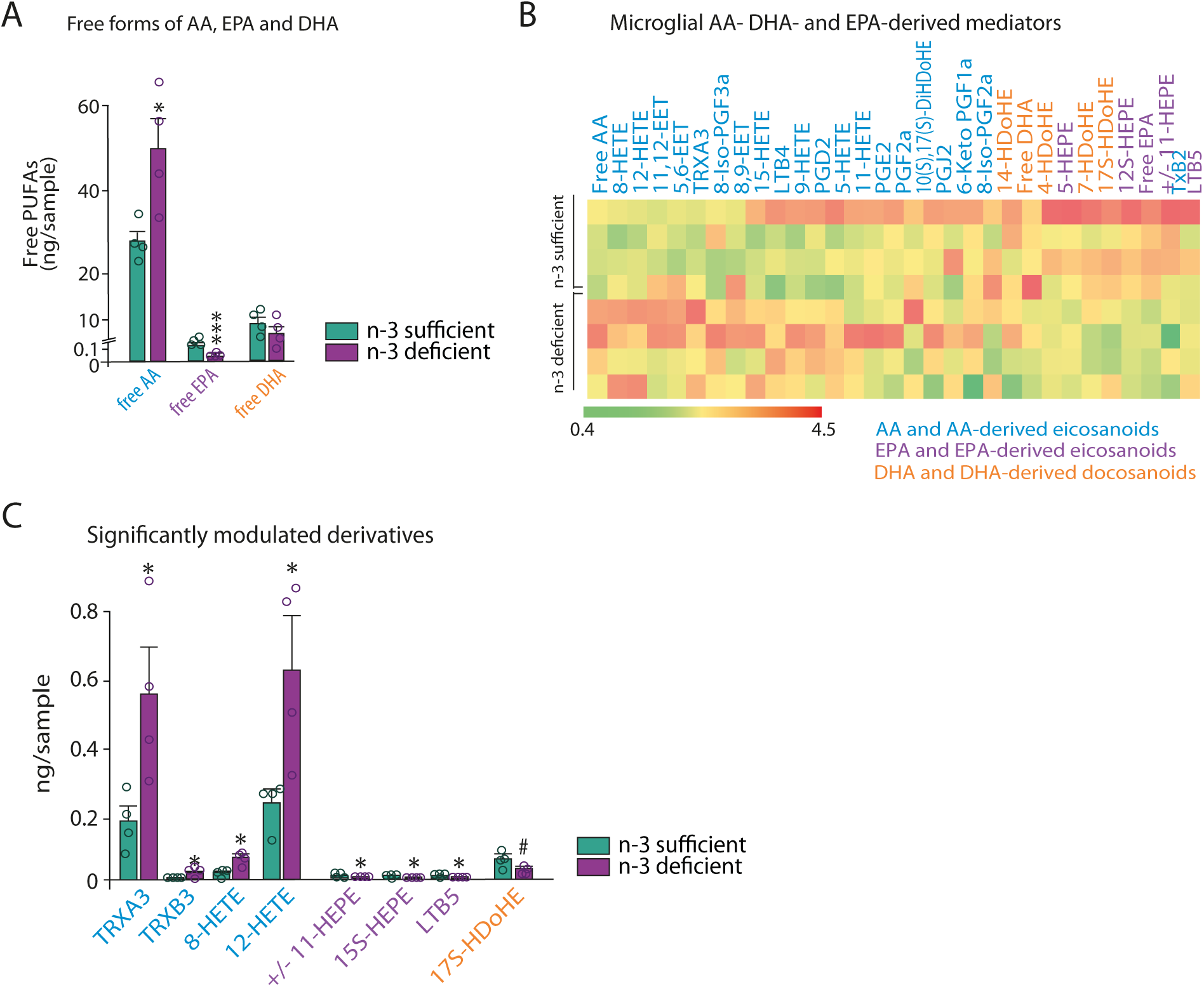
Maternal n-3 PUFA deficiency alters lipid profile in microglia. **A.** Quantification of free (unesterified) forms of AA, EPA and DHA levels in microglia. Means ± SEM; n=4 mice per group. Two-tailed unpaired Student’s t-test; t=3, *p=0.024, free AA; t=6.12, ***p=0.0009, free EPA; t=1.149, p=0.2942, free DHA. **B.** Heat map of all AA-, EPA- and DHA-derived intracellular mediators expressed by microglia. **C.** AA-EPA- and DHA-derived mediators that are significantly modulated by early-life n-3 PUFA deficient diet. Means ± SEM; n=4 mice per group. Two-tailed unpaired Student’s t-test; t=2.693, *p=0.0359, TRXA3; t=2.867, *p=0.0286, TRXB3; t=3.028, *p=0.0231, 8-HETE; t=2.886, *p=0.0278, 12-HETE; t=3.656, *p=0.0106, +/−11-HEPE; t=3, *p=0.024, 15S-HEPE; t=3, *p=0.024, LTB5; t=1.975, *p=0.0957, 17S-HDoHE.

### Activation of a 12/15-LOX/12-HETE signaling pathway increases microglial phagocytic activity under maternal n-3 PUFA deficiency

We continued to dissect the molecular mechanisms linking n-3 PUFA deficiency and microglial phagocytosis of dendritic spines by assessing the oxylipins identified above. We first tested the potential role of AA, EPA and DHA free forms and their bioactive metabolites identified in Figure 6 (except trioxilins TRXA3 and TRXB3, which are not commercially available) on microglial phagocytosis *in vitro*. We applied synaptosomes that were conjugated to a pH-sensitive dye (pHrodo), as a surrogate of synaptic refinement processes on cultured microglia (Figure 7A). Twenty-four hour-application of AA on microglial cells significantly increased their phagocytic activity towards synaptosomes (Figure 7B). These observations confirmed our data showing that AA increased phagocytosis of latex beads by primary microglial cells (Supplementary Figure 7A-B). Conversely, incubation with DHA significantly decreased phagocytosis (Figure 7B), while EPA and DPA n-6 had no effect when compared to controls (Figure 7B). Then, we applied PUFA bioactive mediators for 30 minutes before applying pHrodo synaptosomes and studied the kinetics of phagocytosis over the following 120 minutes. We show that the AA-derived 12-HETE significantly increased microglia phagocytosis of synaptosomes while 8-HETE had no effect (Figure 7C). None of the EPA- and DHA-derived bioactive mediators affected primary microglia phagocytic activity (Figure 7D). For all conditions, the percentage of phagocytic cells is significantly higher at 120 min post-treatment (Figure 7C-D).

**Figure 7:**
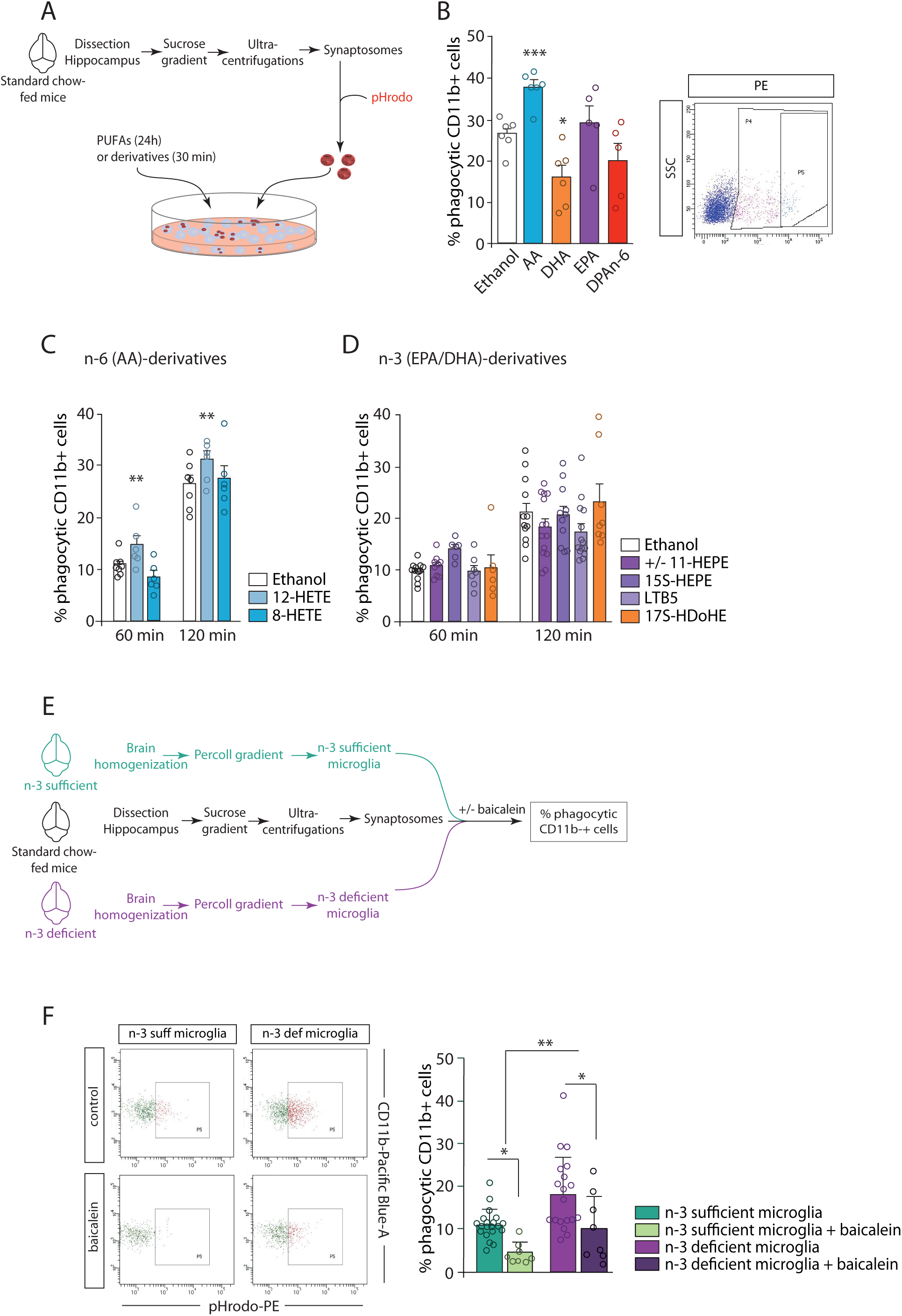
Maternal n-3 PUFA deficiency exacerbates microglial phagocytic activity towards synapses by activating the 12/15-LOX/12-HETE signaling pathway. **A.** Experimental setup. **B.** FACS analysis of phagocytic uptake of pHrodo-labeled synaptosomes by CD11b^+^ microglial cells in primary culture exposed to PUFAs. Means ± SEM; n=5-6 per condition. Two-tailed unpaired Student’s t-test; ***p=0.0005, AA; *p=0.0118, DHA; p=0.55, EPA; p=0.153; DPA n-6. **C-D.** FACS analysis of phagocytic uptake of pHrodo-labeled synaptosomes by CD11b^+^ microglial cells in primary culture exposed to n-6 AA-derived (**C**) or n-3 EPA- and DHA-derived (**D**) lipids. Means ± SEM; n=6-14 per condition. AA derivatives: Two-way ANOVA: time effect, F(1,33)=171.8, ***p<0.0001; treatment effect, F(2,33)=5.538, **p=0.0084; interaction, F(2,33)=0.686, p=0.51. EPA and DHA derivatives: Two-way ANOVA: time effect, F(1,64)=57.25 ***p<0.0001; treatment effect, F(2,64)=1.462, p=0.239; interaction, F(2,64)=1.904, p=0.157. **E.** Experimental setup. **F.** FACS analysis of phagocytic uptake of pHrodo-labeled synaptosomes by freshly sorted n-3 deficient and n-3 sufficient CD11b^+^ microglial cells, exposed to baicalein or its solvent. Two-way ANOVA: microglia lipid status effect, F(1,51)=12.6 ***p=0.0008; treatment effect, F(1,51)=16.06, p=0.0002; interaction, F(1,51)=0.1912, p=0.6638.

AA is metabolized into 12-HETE *via* the 12/15-LOX enzyme and its activity is blocked by baicalein, which reduces 12-HETE production^55–57^. We then tested whether inhibiting 12/15-LOX activity was able to restore normal phagocytic activity in n-3 PUFA deficient microglial cells. We used an *ex vivo* assay in which we exposed freshly sorted n-3 sufficient or n-3 deficient microglia to pHrodo synaptosomes with or without the 12/15-LOX inhibitor baicalein (Figure 7E). We show that baicalein significantly reduced synaptosome phagocytosis by n-3 deficient microglia (Figure 7F). These results confirm that the 12/15-LOX/12-HETE pathway is active in n-3 deficient microglia to increase their phagocytic activity towards synaptic elements.

## DISCUSSION

The phagocytic activity of microglia during development is crucial in shaping neuronal networks, and dysfunction of this activity contributes to neurodevelopmental disorders^26, 58–61^. Here, we identify an unknown function of 12-HETE, a LOX-induced metabolite of AA, in the regulation of microglial phagocytosis. This 12/15-LOX/12-HETE pathway may also contribute to alterations in spine density and connectivity observed in synaptopathies and neurodevelopmental disorders^66^. Moreover, we show that low maternal n-3 PUFA intake-induced modification of microglia lipid composition is the main driver leading to exacerbation of spines phagocytosis in the offspring. This process is complement-dependent, a mechanism that has been repeatedly reported to be the molecular bridge of microglia phagocytosis of spines^25, 49, 62–65^. Our data also reveal that n-3 PUFA deficiency alters the shaping of hippocampal neurons and spatial working memory in pups. Considering that n-3 PUFAs also determine offspring’s microglia homeostatic molecular signature and lipid profile, our study suggests that maternal nutritional environment is crucial for proper microglial activity in the developing brain. These findings further support the importance of n-3 PUFAs in the neurodevelopmental trajectory, pointing to their role in microglia-spine interactions.

Previous clinical and epidemiological studies revealed a relationship between dietary n-3 PUFA content, brain development and the prevalence of neurodevelopmental disorders^17^. The n-3 PUFA index, which reports the level of n-3 PUFAs (EPA+DHA) in erythrocytes and that is used as a biomarker for cardiovascular diseases^67^, is now considered as a new biomarker for several neurodevelopment diseases such as ASD^68^, attention deficit hyperactivity disorder (ADHD)^69^ and schizophrenia^70, 71^. Since these essential fatty acids are necessary for healthy brain development, low dietary supply or impairment of their metabolism is suggested to be involved in the etiology of several psychiatric diseases, including neurodevelopmental disorders^1, 72, 73^. Indeed, decreased DHA levels have been reported in several brain regions of patients diagnosed with such disorders^17, 74–76^. Low n-3 PUFA index is also associated with reduced brain size or connectivity^77, 78^, as well as to reduced cortical functional connectivity during an attentional task^78^. However, until now, the cellular and molecular processes linking low n-3 PUFA levels and altered connectivity in the developing brain were unknown. Our data shed light on an oxylipin-dependent mechanism underlying these clinical observations.

Previous studies conducted in rodents and primates showed that early-life dietary n-3 PUFA deficiency impairs learning and memory at adulthood^79–81^. Here, we show that perinatal n-3 PUFA deficiency-induced memory deficits occur as soon as weaning. This corroborates observations in humans that prenatal and cord blood n-3 PUFA levels are positively correlated with later-life cognitive abilities in infants^82, 83^. In our study, cognitive alterations were associated with a decrease in dendritic length, whereas the complexity of dendritic arborization was unchanged, supporting previous reports^84, 85^. A significant decrease of post-synaptic scaffold proteins PSD-95 and Cofilin in the hippocampus of n-3 deficient mice at weaning, suggests an early alteration of synaptic plasticity^34, 85^. This is in line with a previous study showing that loss of PSD-95 is inversely correlated with DHA dietary supply in a mouse model of Alzheimer’s disease^86^ suggesting that PSD-95 expression depends on DHA levels in a broader context than just neurodevelopment. N-3 PUFA deficiency-altered dendritic arborization has been associated with functional alterations of neuronal networks in the adult hippocampus and cortex^85, 87^ and atypical whole-brain functional community structure^88^, a surrogate of cognitive deficits^89^. Finally, our finding of abnormal hippocampal neuronal spine density in n-3 PUFA deficient mice at weaning corroborates data previously found^84, 90^.

Lipid changes driving metabolic alterations in microglia and their impact on phagocytosis in the developing brain are understudied. Knowing that microglia are now regarded as key controllers of synaptic architecture due to their phagocytic activity^24, 25, 27, 91^, we show for the first time that specific lipid mediators drive microglia phagocytic activity towards synapses. Oxylipins are involved in the regulation of inflammation and phagocytic activity of immune cells^1, 9–11, 54, 95–97^. Their production is largely influenced by n-3 PUFA dietary supply, including in the brain^54^. Using lipidomics, we examined their role in microglial phagocytic activity during development. We show that n-3 PUFA deficient microglia have a unique oxylipin profile, linked to altered free forms of AA, EPA and DHA. AA and its LOX-derived mediator 12-HETE promote microglia phagocytosis of synaptosomes. Conversely, DHA (but not its LOX derivative) decreased microglial phagocytosis. 12-HETE is produced by the 12/15 LOX enzyme^98, 99^ and conversion of AA by 12-HETE is enhanced in n-3 PUFA deficient microglia in the context of brain injury, autoimmune and neurodegenerative diseases^100–102^. 12/15 LOX is also expressed in the developing human brain by CD68-positive microglia and oligodendrocytes^100^. We show that blocking activity of 12/15-LOX by baicalein^56, 57^ attenuates the excessive n-3 PUFA deficient microglia phagocytic activity, which is coherent with the role of this enzyme in mediating phagocytic activity of apoptotic cells by macrophages^10^. Hence, we describe a cell-specific lipid profile of microglia and provide a comprehensive description of the processes involved in n-3 PUFA deficiency-induced microglial alterations, including 12/15-LOX/12-HETE pathway as the molecular trigger enhancing the phagocytic activity of microglia towards spines.

Deficits in microglia-mediated synaptic refinement lead to dysfunctional neuronal networks and behavioral abnormalities resembling some aspects of neurodevelopmental disorders^26, 58–61^. Here, we show that n-3 PUFA deficient microglia display an altered homeostatic molecular profile at weaning with features similar to neurodegenerative microglia, previously reported as more phagocytic^92^. We further identified the complement system, and not PS exposure, as the molecular mechanism driving spine phagocytic activity upon n-3 PUFA deficiency. This is consistent with previous work showing that in the hippocampus, PS-dependent and complement-independent trogocytosis of presynaptic element remodels neuronal network during normal brain development ^28^, while complement-dependent phagocytosis of synapses is promoted in models of brain diseases presenting neuronal network abnormalities, including neurodevelopmental disorders ^17, 24, 25, 47, 93, 94^.

Overall, our data also describe a novel role for the 12/15-LOX/12-HETE oxylipin pathway as a crucial metabolic signaling axis in the regulation of n-3 PUFA deficient microglia activity. We also delineate for the first time that maternal dietary n-3 PUFA deficiency drives microglia in the offspring into a complement-dependent phagocytic phenotype, aberrantly reducing synaptic elements, which drive hippocampal synaptic network dysfunction and spatial working memory deficit. Our findings not only support the necessity of adequate n-3 PUFA status during brain development^103–105^, but also reveal that maternal lipid nutrition, by modulating microglial lipid metabolism and sensing, can be an important determinant of neurodevelopmental synaptopathies such as ASD^66^.

Regarding the clinical implications, these data highlight the relevance of diagnosing DHA/EPA deficits in expectant mothers and in early-life. We provide novel knowledge on highly targetable cellular and molecular pathways linking maternal nutrition and neurodevelopmental disorders. Indeed, specific dietary strategies can enhance PUFA status in microglia^106^, being promising avenues for the prevention and treatment of neurodevelopmental disorders.

## AUTHORS CONTRIBUTION

The data reported in this study can be found in the supplementary materials. C.M., Q.L., L.M., C.L., C.B.B., J.B., A.T., J.C.D., A.S(ere)., A.A. and V.D.P. performed most experiments. K.H. performed and R.P.B. oversaw the lipid analyses of microglia. S.B. and A.S(ierra). performed and A.S. oversaw the apoptosis experiments. C.L., K.B. and M.E.T. performed and M.E.T. oversaw the E.M. experiments. S.G. and L.B. performed and analyzed lipid experiments on whole hippocampus. J.B. performed and analyzed BBB experiments. C.M. performed and O.B. oversaw transcriptomic analyses. A.D.G. analyzed transcriptomic data. N.J.G. performed FACS phagocytosis experiments. L.F. performed and analyzed TAM receptors Western blots. P.G. contributed to the design of the experiments. C.J. performed whole structure lipid composition experiments and conducted the data analyses. S.L. and A.N. equally supervised the entire project and wrote the manuscript. All authors proof-read the manuscript.

## ACKNOWLEDGMENTS

Funding for this research was provided by the Institut National pour la Recherche Agronomique (INRA), the Bordeaux Univ, the Foundation for Medical Research (FRM) (DEQ20170336724), the French Foundation (FDF, #00070700). CM was funded by the French Ministry for Research and Higher Education, CBB was supported by the French National Agency for Research (ANR-12-BSV4-0025-04) and Agreenskill, JCD was supported by the Nouvelle Region Aquitaine (2011.1303003 POST-DOCTORAT), QL was supported by the Region Ile de France (PICRI, the Ceberal Palsy Foundation #13020605), AT was supported by the French National Agency for Research (ANR-2010-BLAN-141403), ADG was supported by Agreenskill and Marie-Curie European Grant, CL was supported by Idex grant. AS was supported by grants from the Spanish Ministry of Economy and Competitiveness with FEDER funds to AS (BFU2015-66689, RYC-2013-12817), a BBVA Foundation Grant for Researchers and Cultural Creators to AS, and a Basque Government grant (PI_2016_1_0011) to AS. PG was supported by Inserm, Université de Paris, Investissement d’Avenir (ANR-11-INBS-0011, NeurATRIS), ERA-NET Neuron (Micromet). MET, CL and KB were supported by grants from NARSAD and the CIHR awarded to MET.

We thank R. Van Der Wickt and Celine Lucas for technical assistance. This work benefited from the facilities and expertise of the imaging platform Imag’In (www.incia.u-bordeaux1.fr), which is supported by CNRS and Region Aquitaine and from the Bordeaux Imaging Center. We thank Atika Zouine and Vincent Pitard for technical assistance at the Flow cytometry facility, CNRS UMS 3427, INSERM US 005, Univ. Bordeaux, F-33000 Bordeaux, France. Analysis of eicosanoids and docosanoids was performed at the Analytical Facility for Bioactive Molecules (AFBM) of the Centre for the Study of Complex Childhood Diseases (CSCCD) at the Hospital for Sick Children, Toronto, Ontario. CSCCD was supported by the Canadian Foundation for Innovation (CFI). The Western blot analyses and synaptosome extractions were done in the Biochemistry and Biophysics Platform of the Bordeaux Neurocampus at the Bordeaux University funded by the LABEX BRAIN (ANR-10-LABX-43) with the help of Y.Rufin.

---

**Supplementary Figure 1.**
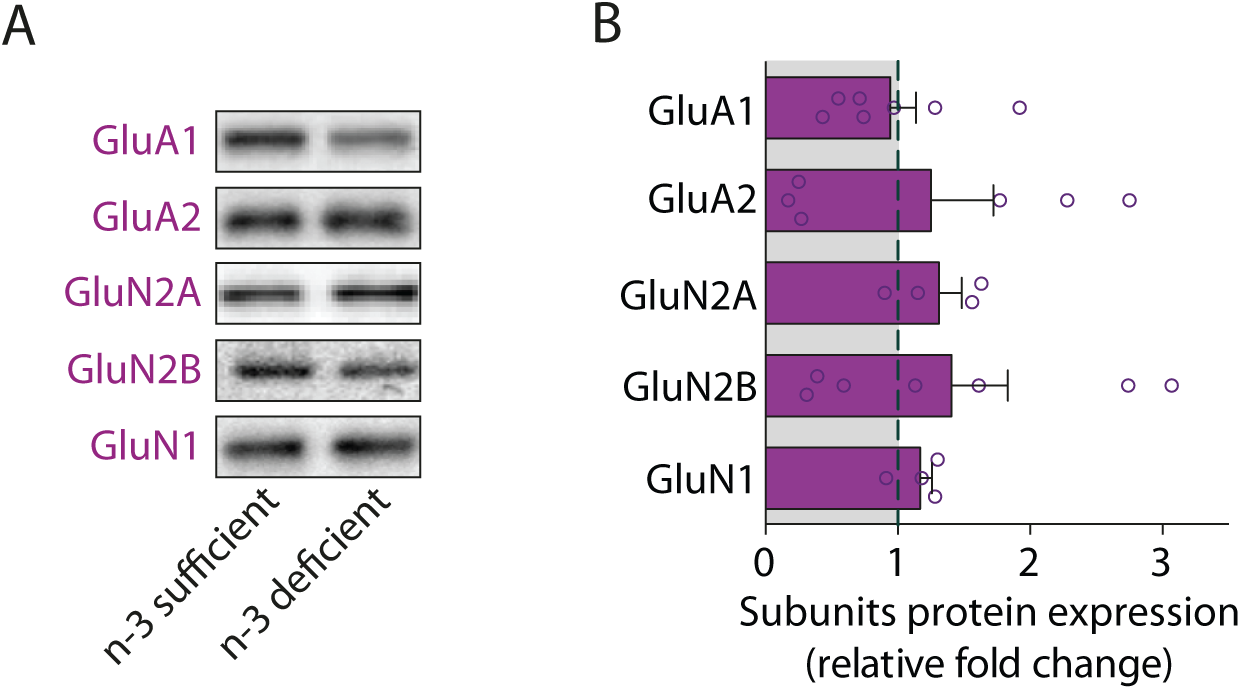

**Supplementary Figure 2.**
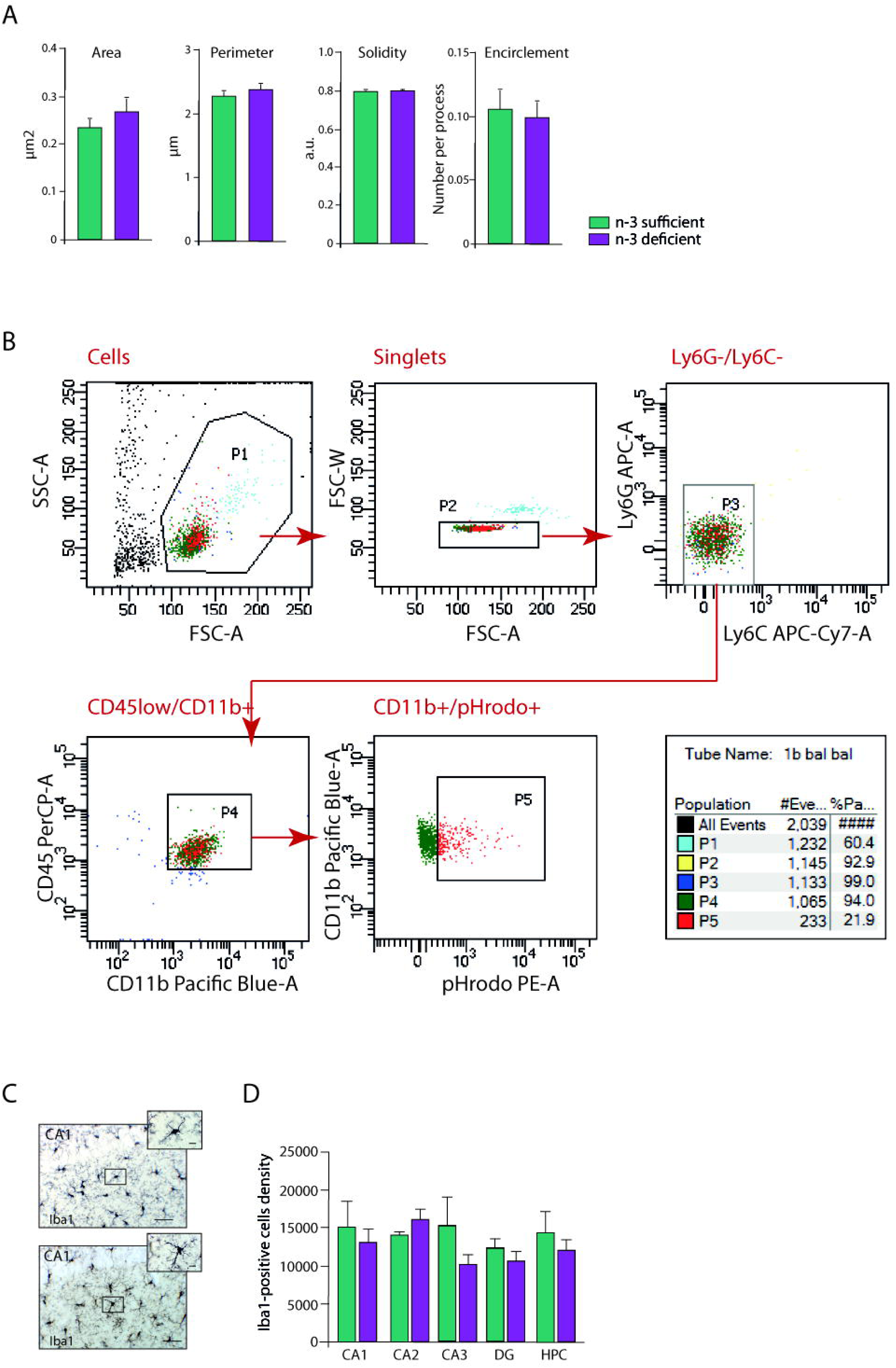

**Supplementary Figure 3.**
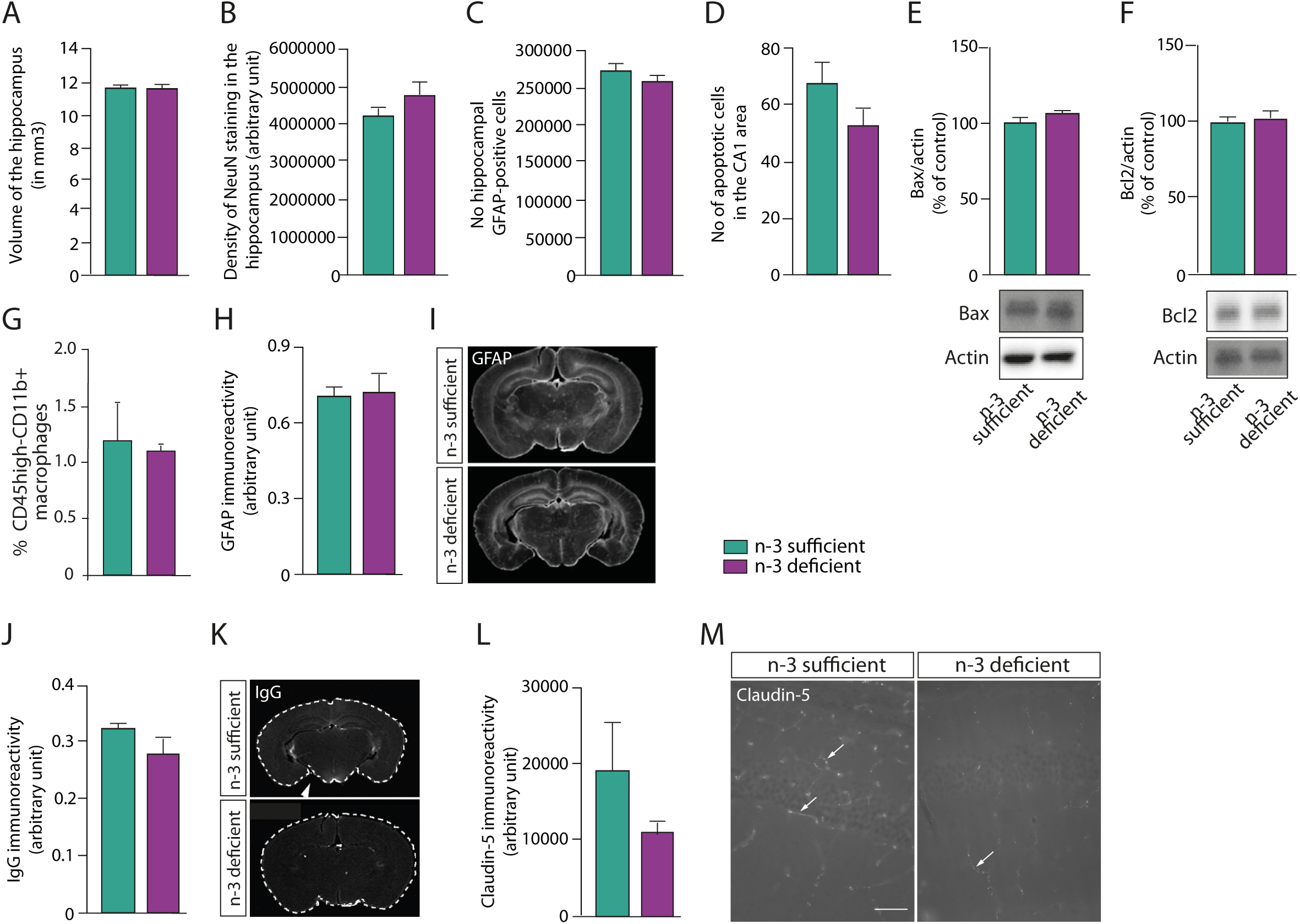

**Supplementary Figure 4.**
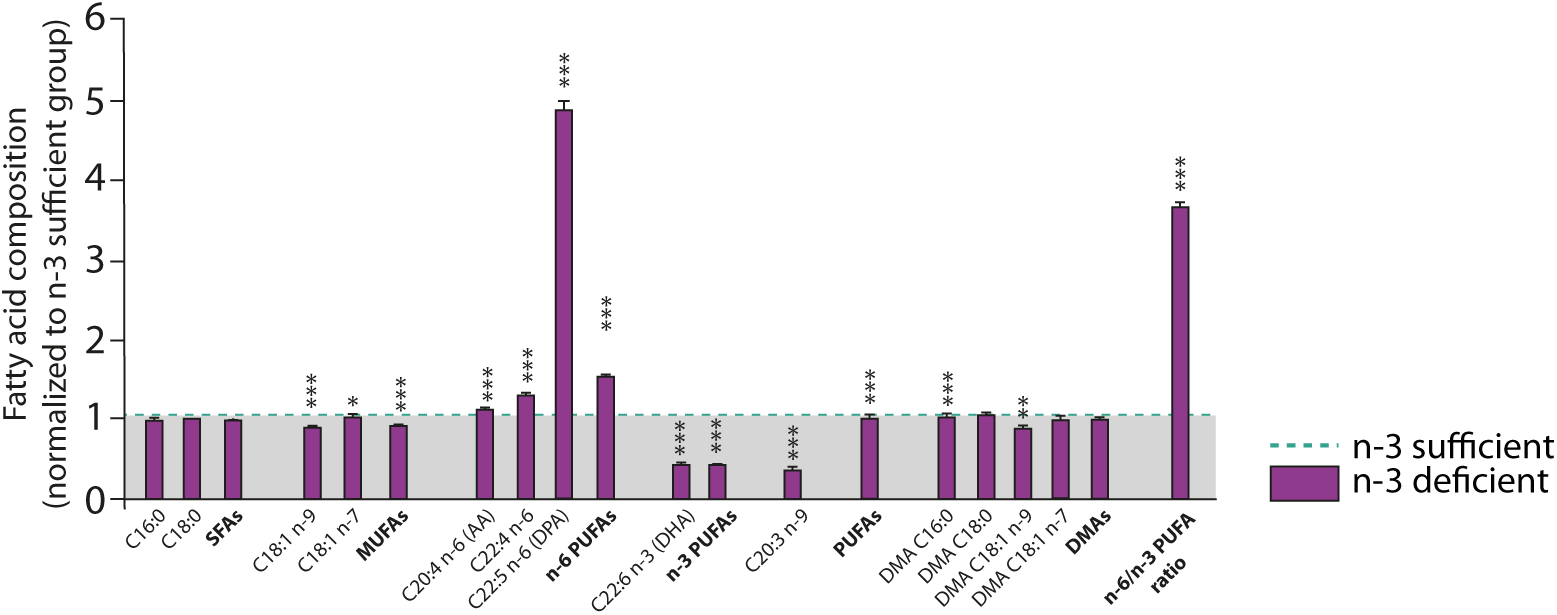

**Supplementary Figure 5.**
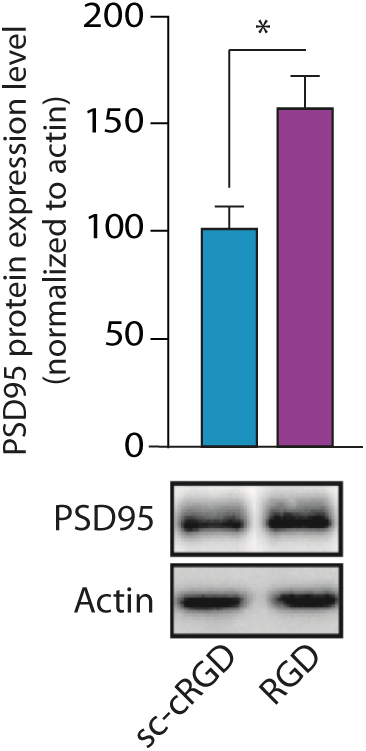

**Supplementary Figure 6.**
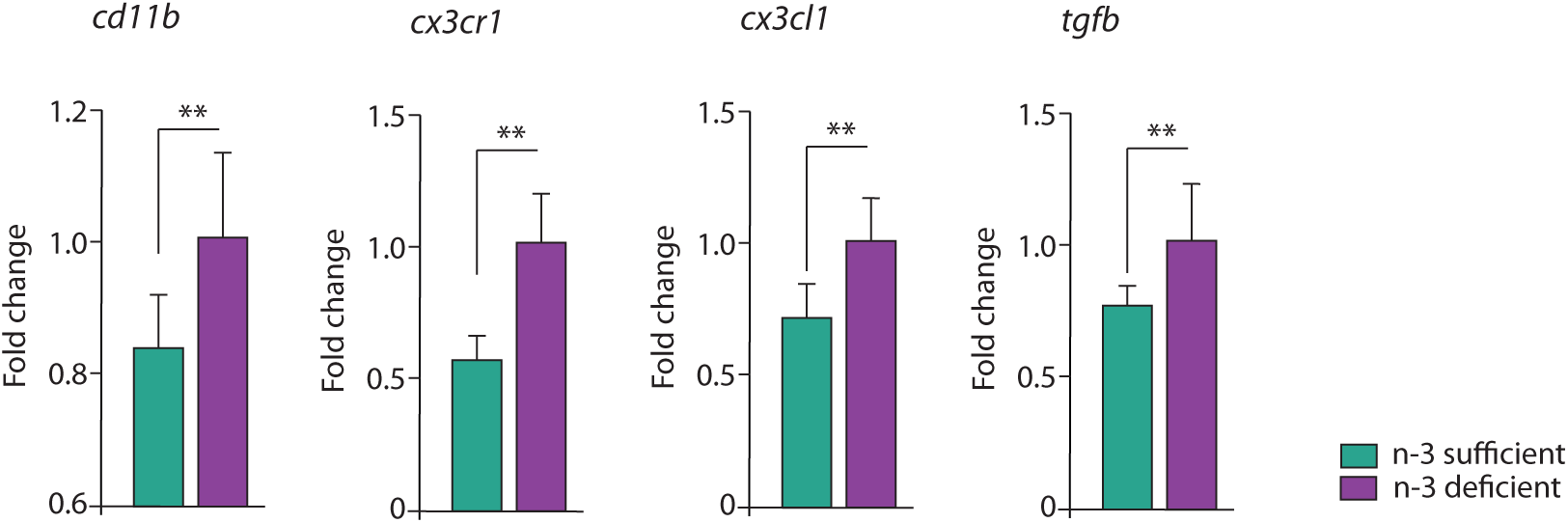

**Supplementary Figure 7.**
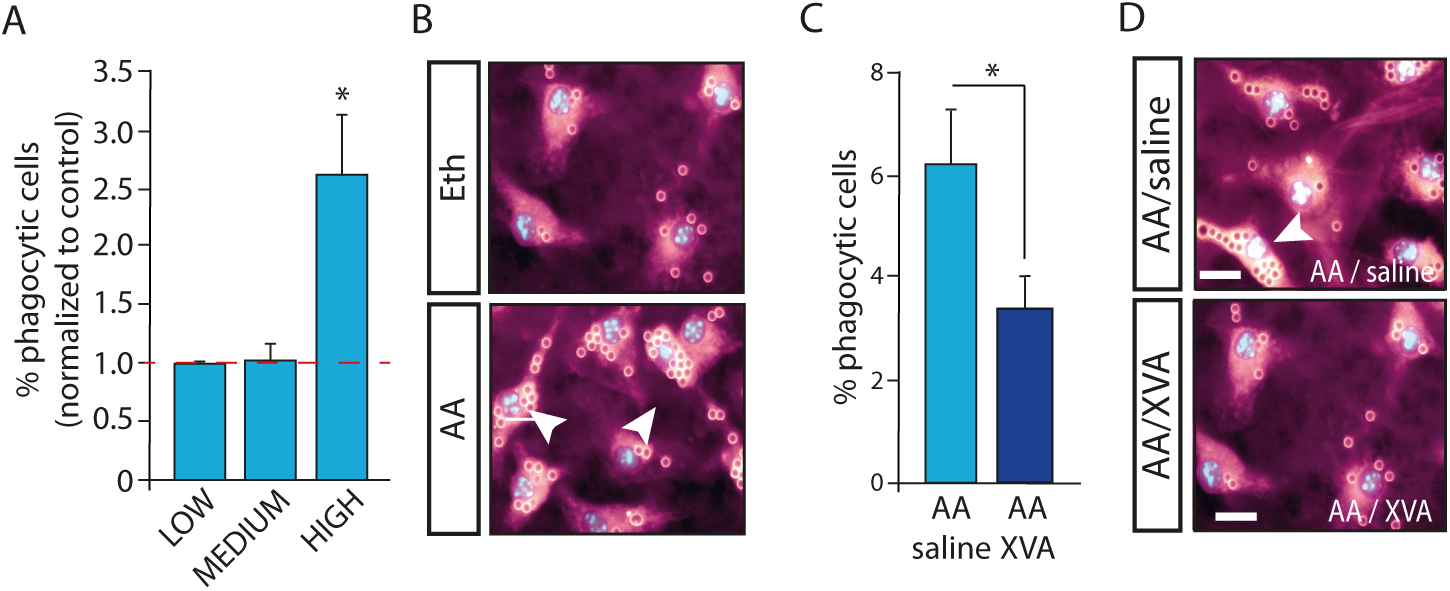

