## Supplementary material for "Essential omega-3 fatty acids tune microglial phagocytosis of synaptic elements in the developing brain": Material and methods

**Animals**

Animal experiments were carried out according to the Quality Reference System of INRA and approved by the local ethical committee for care and use of animals (#APAFIS 4198). Male and female CD1 mice 8-week old were purchased from Janvier Labs (Le Gesnest St Isle, France) and were SPF-housed in temperature and humidity-controlled cages on a 12 h light/dark cycle with food and water *ad libitum*.

**Diet**

After mating, CD1 females were randomly assigned to either a n-3 deficient or a n-3 sufficient diet, both containing 5% fat, in the form of sunflower oil (rich in LA, the 'n-3 deficient diet') or a mixture of different oils of which rapeseed oil (rich in ALA, the 'n-3 sufficient diet') throughout gestation and lactation^1–4^ (Table 1 and 2). All experiments were performed on the offspring from n-3 deficient and n-3 sufficient dams at post-natal day (P) 21.

**Drugs**

Cyclo(Arg-Gly-Asp-D-Phe-Val) (cRGD; Bachem, H-2574) and its control, scrambled, peptide c(RADfV) (or sc-cRGD; Bachem, H-4088) were dissolved in acetic acid (1N) and further diluted into aCSF to a final concentration of 1mM. A volume of 1µl of cRGD or sc-cRGD was injected in the CA1 region of the hippocampus, bilaterally, at P14. Animals were euthanized at P21 and processed for Western blot analysis.

XVA-143 (kindly provided by Hoffmann-La Roche) was dissolved in aCSF at 5mM. 1µl was injected bilaterally in the CA1 region of the hippocampus at P17. Animals were used at P21 for behavioral and biochemical analyses. For primary microglial cell cultures, XVA-143 was applied 30 min before beads incubation at a final concentration of 1µM.

**Cortex fatty acid composition**

Fatty acid composition was studied in the cortex (not hippocampus) of animals as: 1) cortex and hippocampus lipid composition vary similarly under the diets used in this study (Joffre et al., 2016); 2) to reduce the number of animals. Cortical lipids were extracted according to the method of Folch ^5^, fatty acids were transmethylated according to the method of Morrison and Smith^6^ and fatty acid methyl esters (FAMEs) were analyzed on a FOCUS GC gas chromatograph (Thermo Electron Corporation) equipped with a split injector and a flame ionization detector as previously described ^1–4,7^.

**Sorting of microglial cells.**

Brains were homogenized in Hanks’ Balanced Salt Solution (HBSS), pH 7.4 passing through a 70 μm nylon cell strainer. Homogenates were centrifuged at 600g for 6 min. Supernatants were removed and cell pellets were re-suspended in 70% isotonic Percoll (GE-Healthcare, Aulnay sous Bois, France). A discontinuous Percoll density gradient was set up as follows: 70%, 35% and 0% isotonic Percoll. Gradients were centrifuged at 2000g for 20min. Microglia cells were collected at the interphase between the 70% and 35% Percoll layers. Cells were washed and counted with a hemacytometer. For each brain extraction approximately 3×10^5^ cells were isolated ^8^.

**Microglia lipidomics**

***Fatty acids.*** Extraction and quantification of total fatty acids by GC-FID: Microglia from 3-4 brains per sample were lysed in 100% methanol and extracted via from the Folch method ^5^ (2:1:0.8 chloroform: methanol: 0.88% KCl) and analyzed by a Varian-430 gas chromatograph (Varian, Lake Forest CA) as described previously ^9,10^.

***Lipid mediators.*** Isolation and quantification of brain eicosanoids and docosanoids by LC-MS-MS: Pooled microglia extracted from 3-4 brains per sample were lysed in 15% methanol and stored at -80°C prior to analysis. One ng of internal standard mixture was added to each sample, and lipid mediators were extracted, analyzed by LC-MS-MS and quantified as previously described ^10^.

**RNA isolation and Nanostring RNA counting**

Total RNA was extracted from isolated microglia using mirVanaTM miRNA isolation kit (Ambion) according to the manufacturer's protocol. We performed nCounter multiplexed target profiling of 542 microglial transcripts (MG550). MG550 encompasses 400 unique and enriched microglial genes we have identified previously ^11^ and additional 150 inflammation-, inflammasome- and phagocytosis-related genes. 100 ng of total RNA per sample were used in all described nCounter analyses according to the manufacturer's protocol ^11^.

**Network Analysis**

To investigate gene coexpression relationships between groups, a pairwise transcript-to-transcript matrix was calculated in the software tool Miru (Kajeka, UK) from the set of differentially expressed transcripts using a Pearson correlation threshold r= 0.85. A network graph was generated where nodes represent individual probe sets (transcripts/genes), and edges between them correlation of expression pattern with Pearson correlation coefficients above the selected threshold. The graph was clustered into discrete 6 groups of transcripts sharing similar expression profiles using the Markov clustering algorithm (inflation 2.2).

**Electron microscopy**

Mice were anesthetized with sodium pentobarbital (80 mg/kg, i.p.) and perfused with 0.2% glutaraldehyde in 4% PFA. Transverse sections of the brain (50µm thick) were cut in PBS (50 mM at pH 7.4) using a vibratome and stored at -20^o^C in cryoprotectant solution (30% glycerol and 30% ethylene glycol in PBS) ^12^.

Sections were washed in PBS and quenched with 0.3% H_2_O_2_ in PBS for 5 min and then with 0.1% NaBH_4_ for 30 min at room temperature (RT), washed in Tris–buffered saline (TBS; 50 mM at pH 7.4) and processed freely-floating for immunoperoxidase staining. Sections were pre-incubated for 1h at RT in a blocking solution of TBS containing 10% fetal bovine serum, 3% bovine serum albumin, and 0.01% Triton X100, before overnight incubation at 4^o^C in rabbit anti-Iba1 antibody (1:1000; Wako Pure Chemical Industries) and rinsed in TBS. After incubation for 1.5h at RT in goat anti-rabbit IgGs conjugated to biotin (1:200 in blocking solution; Jackson Immunoresearch) and for 1 h with ABC Vectastain mix (1:100 in TBS; Vector Laboratories), the labeling was revealed using diaminobenzidine (DAB; 0.05%) and hydrogen peroxide (0.015%) in TBS. After immunostaining, sections were post-fixed flat in 1% osmium tetroxide and dehydrated in ascending concentrations of ethanol. They were treated with propylene oxide, impregnated in Durcupan (EMS) overnight at RT, mounted between ACLAR embedding films (EMS), and cured at 55^o^C for 72 h. Areas of CA1 stratum radiatum were excised from the embedding films and re-embedded at the tip of resin blocks. Ultrathin (65–80 nm) sections were cut with an ultramicrotome (Leica Ultracut UC7), collected on bare square-mesh grids, and examined at 80 kV with a FEI Tecnai Spirit G2 transmission electron microscope.

Pictures were randomly taken at 9300X in the CA1 stratum radiatum of each animal, for a total surface of ~2000 µm2 of neuropil captured per animal, using an ORCA-HR digital camera (10 MP; Hamamatsu). Cellular profiles were identified according to criteria previously defined ^12–14^. For quantitative analysis, each captured Iba1-positive microglial process was analyzed. A phagocytic index was compiled by summing up the vacuoles and endosomes containing cellular materials such as membranes, axon terminals with 40-nm synaptic vesicles and dendritic spines with a postsynaptic density, on a microglial process basis ^15^.

**Immunohistochemistry**

Mice were deeply anesthetized with isoflurane and transcardially perfused with PBS followed by 4% PFA. Brain was removed, post-fixed in PFA overnight at 4°C and cryoprotected in 30% sucrose at 4°C. Immunohistochemistry experiments were performed on free-floating coronal 30 μm cryostat slices. The following antibodies were used: 1:1,000 rabbit anti-Iba1 (Wako, **#019-19741**), 1:100 mouse anti-PSD95 (Cell Signaling Technology, ABIN1304920), 1:500 rat anti-CD11b (**AbD Serotec, #MCA711**), 1:500 rabbit anti-C1q (**Abcam, #ab182451**), 1:20 mouse anti-C3aR (**Hycult biotech, #HM1123**), 1:1000 rabbit anti-GFAP (Dako, Z03334), 1:1000 mouse anti-NeuN (**Millipore, #MAB377**),1:200 rabbit anti-Annexin V (Abcam, #ab14196), 1:500 mouse anti-claudin 5 (Life Technologies: Invitrogen), 1:500 IRDye 800 conjugated affinity purified goat-anti-mouse IgG (Rockland, Gilbertsville, PA). Primary antibodies were visualized with appropriate secondary antibodies conjugated with Alexa fluorophores (Invitrogen) and counterstained with DAPI or with biotin. When biotinylated, secondary antibodies were revealed using the streptavidin-biotin-immunoperoxidase technique, giving a black precipitate.

**Image analysis**

*Densitometry*. Individual images were analyzed with Fiji or Image J software (Image J, open source), using the following procedure: 1) user-defined thresholding value applied to each image, 2) calculation of area of staining from background for each protein of interest. Final values are represented as a surface area in pixel values. Control sections for all studies in which primary or secondary antibodies were omitted resulted in negative staining (not shown).

*Stereological analysis*. Iba1- and GFAP-immunoreactive cell numbers and volume of the hippocampus were thoroughly determined in the hippocampus with the unbiased stereological sampling method based on optical dissector stereological probe, as previously described ^2^.

*Apoptotic cells counting*. The number of apoptotic cells (cells with pycnotic/karryorhectic morphology) was estimated using unbiased stereology methods^16^, and is reported as cells/mm3.

**Cell culture and treatments**

For *in vitro* phagocytic assays, primary mixed glial cell cultures were prepared from the cortices of P0–1 CD1 mice. After dissection in 0.1M PBS with 6% glucose and 2% penicillin–streptomycin (Gibco, Cergy Pontoise, France) and removal of the meninges, the cortices were chopped and subsequently mechanically dissociated. The suspension was diluted in low glucose DMEM (31885, Gibco) supplemented with 10% fetal bovine serum (Gibco) and 1% penicillin–streptomycin. Microglia were isolated from primary mixed glial cultures on DIV14 using a reciprocating shaker (45 min at RT) and repeated rinsing with their medium using a 10 mL pipette. Media was subsequently removed, microglia pelleted via centrifugation (2000 rpm × 10 min) and following resuspension maintained in DMEM supplemented with 10% FBS at a concentration of 4 × 10^5^ cells/mL in 6-well culture plates. Cells were then treated for 24 h in serum-free medium containing 50 µmol/L fatty acid-free bovine serum albumin (BSA) added either with DHA, AA, DPAn-6 or EPA 30 µmol/L in 0.1% ethanol or 0.1% ethanol. For oxylipins, 30min prior to the phagocytosis assay, culture medium was replaced by serum-free DMEM containing 100nM of the given lipid in 0.03% ethanol, or 0,03% ethanol.

**Phagocytic assay**

*Ex vivo* assays: the phagocytic capacity of microglia was determined by the level of pHrodo™ Red fluorescence accumulated in the cells. First we used pHrodo-conjugated E*.Coli* bioparticles (adapted from^17^). A suspension of 2.5x105 sorted microglia cells in 100 μL of RPMI/1% BSA was incubated with bioparticles in an incubator at 37°C and 5% of CO_2_ for each time point. Cells were then pelleted by centrifugation at 1000g for 10 min at 4°C, resuspended in 200 μL of RPMI/1 %BSA and stained with CD11b-APC antibody for quantification by cytometry. We also developed a model using pHrodo-conjugated synaptosomes as a substrate for the phagocytosis, adapted from ^18^. Synaptosomes were prepared from P21 mice hippocampus as described in ^19^, and labeled with pHrodo Red, succinimidyl ester (Thermo Fisher Scientific, P36600) according to manufacturer instructions. Synaptosomes were resuspended at a concentration of 20 mg/ml in DMEM. Microglia were plated at a density of 5.10^5 cells/well in 24-well plates and received 300 μg of synaptosomes for 120 min in an incubator at 37°C and 5% CO_2._ When needed, baïcalein was applied at a concentration of 10 μM 90 min prior to the beginning of the essay (i.e. the addition of synaptosomes). Cells were collected by gentle trypsin treatment, rinsed and resuspended in PBS/1%BSA buffer to be stained with CD11b-V450 and CD45 Cy5 for analysis by cytometry. After selection of the CD11b+/CD45low, cells were gated on PE channel for quantification of pHrodo fluorescence.

Experiments in Figure 3C and Figure 7D have been performed on the same batches in order to spare animals. Hence, data used for Figure 3C control groups are a randomly chosen subset of figure 7D’s data.

*In vitro,* two different substrates where used: FCS-coated Yelloworange fluorescent carboxylated microspheres (Fluoresbrite® YO Carboxylate Microspheres 3.00µm***,*** Polysciences Europe GmbH, #19393-5) or the pHrodo-conjugated synaptosomes as described above. To quantify their phagocytic index, primary microglial cell cultures were incubated with a suspension of microspheres at a concentration of 1.1×10^7^ microspheres/ml for 30 min at 37°C, intensively washed and finally fixed with 4% paraformaldehyde. Cells were stained with anti-Iba1 antibodies. A blinded experimenter counted beads per cell. We calculated the phagocytic index as in^20^. To measure phagocytic activity by cytometry, cells were incubated for 60 or 120 min with 300 μg synaptosomes per well. Medium was removed and cell collected following trypsin treatment, pelleted, rinsed in PBS/1%BSA and stained with CD11b-FITC antibody for cytometry analysis.

**Quantitative real-time PCR**

Total RNA was extracted from hippocampi using TRIzol (Invitrogen, Life TechnologiesTM). RNA purity and concentration were determined using a Nanodrop spectrophotometer (Nanodrop technologies, Wilmington, DE). 2μg of RNA was reverse transcribed to synthesize cDNA using Superscript III (Invitrogen, Life TechnologiesTM) and random primer according to the manufacturer’s protocol. Quantitative PCR were performed on 384-well plates using epMotion 5070 (Eppendorf). 10µl of cDNA diluted 1:5 (20ng/µl) were amplified by real-time PCR. Primer references: C1qa Mm00432142_m1; C3 Mm00437838_m1; TREM2 Mm00451744_m1; CD11b Mm01271262_m1 (for in vitro experiments); CD33 Mm00491152_m1; CX3CR1 Mm00438354_m1; CX3CL1 Mm00436454_m1; TGFb Mm03024053_m1; beta-2 microglobulin Mm00437762_m1 (housekeeping gene) (Life Technologie). For *in vivo* quantification of CD11b mRNA expression, we used SYBR Green technology. Ten µl of cDNA diluted 1/60 (1,66ng/µL) were amplified by real-time PCR. Primers sequences: CD11b Forward AATGATGCTTACCTGGGTTAT GCT/Reverse TGA TAC CGA GGT GCTCCTAAAAC; Housekeeping gene beta-2 microglobulin: Forward CTGATACATACGCCTGCAGAGTTAA/Reverse GATCACATGTCTCGATCCCAGTAG. For all experiments, the difference between target and housekeeping gene Ct values (ΔCt) was calculated to normalize for differences in the amount of total nucleic acid added to each reaction and in the efficiency of the RT step. The expression of target gene (linear value) normalized to the housekeeping gene was determined by 2^-(ΔCt)^.

**Golgi staining**

Brains were processed for Golgi-Cox staining using the FD Rapid Golgi Staining kit (FD Neurotechnologies, Inc) as described previously ^21^. Brains were left in the staining solution for 12 days, frozen in isopentane solution and kept at -80ºC until sectioning and coloration. The brains were cryostat-cut at a thickness of 100 μm. Sections were mounted on gelatin-coated slides and analyzed using a motorized Leica DM5000 microscope at 63x magnification. The images were acquired using a CCD Coolsnap camera and Metamorph software. For analysis, we randomly selected pyramidal neurons from the CA1 region of the dorsal hippocampus that were fully penetrated by the Golgi coloration and distinguishable from other neurons (n-3 deficient mice had less usable neurons than n-3 sufficient mice).

**Western blotting**

Brains were carefully placed on a glass plate over dry ice to collect hippocampi that were immediately frozen and stored at - 80°C until use. Samples were homogenized in lysis buffer plus anti-phosphatase solution (Tris/HCl 20 mM pH 7.4 with EDTA 1 mM, MgCl2 5 mM, dithiothreitol 1 mM, Na orthovanadate 2 mM, protease inhibitors cocktail 1X and Na fluoride 1 mM). Homogenates were centrifuged 10 min at 5000 rpm to remove nuclei. Supernatants were stored at -80°C. Protein contents were determined by Bio-Rad protein assay according to the manufacturer’s protocol (Bio-Rad) and then heated to 100°C for 5 min in Laemmli sample buffer (2% sodium dodecyl sulfate and 5% dithiothreitol).

Equal quantities of proteins (20μg/well) were electrophoresed onto an 8% or 12% polyacrylamide gel with a 4% stacking gel. Proteins were blotted on PVDF membranes (Immobilon, Millipore, Paris, France). Membranes were saturated by incubation with 5% milk in TBS and Tween 0.1% for 1h. Blots were probed overnight at 4 °C with 1:1,000 rabbit anti-PSD95 (Cell signaling), 1:500 rabbit anti-Bax (Santa cruz, SC-493), 1:500 rabbit anti-Bcl2 (Santa Cruz Biotechnology, SC-492), 1:1,000 mouse anti-Mer (R&D systems, AF591), 1:1,000 mouse anti-Axl (R&D systems, AF854), 1:1,000 rabbit anti-ERK1/2 (Cell Signaling Technology, 137F5), 1:500 goat anti-MFG-E8 (R&D systems, AF2805), 1:1000 rabbit anti-GluA1 (Santa Cruz Biotechnology, SC-28799), 1:1000 rabbit anti-GluA2 (Santa Cruz Biotechnology, SC-7611), 1:1000 goat anti-GluN2A (Santa Cruz Biotechnology, SC-1468), 1:1000 goat anti-GluN2B (Santa Cruz Biotechnology, SC-1469), 1:1000 rabbit anti-GluN1 (Santa Cruz Biotechnology, SC-31556), 1:5000 rabbit anti-SAP102 (Synaptic System, 124213), 1:300 mouse anti-cofilin (Abcam, Ab-54532), 1:5000 rabbit anti-actin (Cell Signaling Technology, 4967), 1:5000 rabbit anti-GAPDH (Cell Signaling Technology, 2118S). After washing in TBS-Tween, membranes were incubated with the secondary antibody coupled to Horse Radish Peroxidase (HRP, Southern Biotechnology Associates, Birmingham, AL, USA) diluted in TBS-Tween supplemented with 3% milk, for 2h at room temperature. Membranes were washed and the complex was detected with an ECL kit (ElectroChemoLuminescence, Amersham, Orsay, France). Optical density capture of the signal obtained was performed with the Syngene Chemigenius2 apparatus (Synoptics, Cambridge, UK). Intensity of the signal was quantified using GeneTools software (Synoptics, Cambridge, UK).

**C3 ELISA assay**

The level of C3 protein located on synaptosomes was measured using the Mouse C3 (Complement Factor 3) ELISA Kit (Genway Biotech GWB-7555C7). Synaptosomes were prepared as described previously ^19^, using the hippocampi of 2 mice per sample. After extraction, synaptosomes were lysed using sonication in TB. Quantifications were performed according to manufacturer instructions, in duplicates and by loading 5 μg of proteins extract per well.

**Behavioral measurements.**

Mice were handled daily and weighed before and during behavioral experiments. Sessions were recorded with a ceiling-mounted video camera and analyzed using Smart software (Panlab, Barcelona, Spain). The Y-maze was used to assess spatial working memory as previously described ^4,7,21,22^. Each arm was 34 cm long, 8 cm wide and 14 cm high. The floor of the maze was covered with corn cob litter which was mixed between each trial to remove olfactory cues. Visual cues were placed in the testing room and kept constant during the whole test. In the first trial, one arm was closed with a guillotine door and mice were allowed to visit two arms of the maze for 5 min. After a 30-min inter-trial interval (ITI), mice were placed back in the start arm and allowed free access to the three arms for 5 min. Start and closed arms were randomly assigned for each mouse. Data are presented as the time spent exploring the novel and the familiar arms during the second trial.

**Statistical analyses**

For most experiments, experimental groups were compared using Student t-test or non-parametric Mann-Whitney test (when equality of variance or normality failed). A two-way ANOVA test was used for Y-maze experiments (n-3 deficient *vs* n-3 sufficient mice), XVA-143 and cRGD experiments (PSD95 Western blotting data), *ex vivo* and *in vitro* phagocytic activity. One sample t-test was used for behavioral analyses of XVA-143 effects. All data were expressed as means ± standard error of the mean (SEM). A p<0.05 was considered as statistically significant.
