## Supplementary material for "Essential omega-3 fatty acids tune microglial phagocytosis of synaptic elements in the developing brain"

**Supplementary Figure 1: Effect of maternal n-3 PUFA deficiency on ionotropic glutamatergic receptor subunits expression level.** **A.** Representative Western blots for GluA1, GluA2, GluN2A, GluN2B and GluN1 subunits. **B.** Quantification of ionotropic glutamatergic receptor protein expression in n-3 deficient mice relative to n-3 sufficient mice. Means ± SEM; n=4-7 mice per group. Two-tailed unpaired Student's t-test, t=1.226, p=0.2597, GluA1; t=0.7894, p=0.4452, GluA2; t=1.413, p=0.201, GluN2A; t=0.417, p=0.6864, GluN2B; t=0.2475, p=0.8087, GluN1.

**Supplementary Figure 2: Maternal n-3 PUFA deficiency does not modify microglial fine morphology and density. A.** EM analysis of the fine morphology of Iba1 positive microglial processes (area, perimeter, solidity i.e. spatial density, and encirclement i.e. ability to enwrap synapses). Means ± SEM; n=627-270 processes per group. Two-tailed unpaired Student's t-test, t=0.9621, p=0.3362, area; t=0.82, p=0.412, perimeter; t=0.487, p=0.6264, solidity; t=0.3607, p=0.7184, encirclement. **B.** Gating strategy for ex vivo microglia phagocytosis experiments corresponding to Figure 2, 3 and 7. Figure shows representative FACS plots of gates P1 (Cells), P2 (Single cells), P3 (Ly6G-ve/Ly6G-ve to gate out neutrophil and monocyte populations), P4 (CD45low/CD11b+ve Microglia), P5 (pHrodo +ve synaptosomes phagocytosed by microglia). **C.** Representative images of Iba1 immunostained microglial cells in the CA1 region of the hippocampus. Scale bar=100µm. Scale bar insert=10µm. **D.** Stereological counting of microglial cells density in the hippocampus of n-3 deficient and n-3 sufficient mice. Means ± SEM; n=3 mice per group. Two-tailed unpaired Student's t-test, t=0.47, p=0.66, CA1; t=1.85, p=0.14, CA2; t=1.34, p=0.25, CA3; t=0.97, p=0.38, DG; t=0.77, p=0.48, whole hippocampus.

**Supplementary Figure 3:** **Maternal n-3 PUFA deficiency does not modify hippocampus volume and cell content, apoptosis, macrophages infiltration and BBB integrity.** **A.** Stereological measurements of hippocampal volume in n-3 deficient and n-3 sufficient mice. Means ± SEM; n=4 mice per group. Two-tailed unpaired Student's t-test, t=1.012, p=0.35. **B-C.** Quantification of NeuN immunostaining density (B) and number of GFAP-positive cells (**C**) within the hippocampus of n-3 sufficient and n-3 deficient mice cells. Means ± SEM; n=4 mice per group. Two-tailed unpaired Student's t-test, p=0.36, NeuN; t=1.64, p=0.15, GFAP. **D.** Quantification of the number of apoptotic cells in the CA1 region of the hippocampus. Means ± SEM; n=7 mice per group. Two-tailed unpaired Student's t-test, t=1.542, p=0.15. **E-F.** Western blot analysis of the pro-apoptotic Bax (**E**) and anti-apoptotic Bcl2 (**F**) protein expression. Means ± SEM; n=6-8 mice per group. Two-tailed unpaired Student's t-test, t=1.515, p=0.155, Bax; t=0.39, p=0.705, Bcl2. **G.** FACS quantification CD45 high-CD11b+ cells from the CNS of n-3 deficient and n-3 sufficient mice. Means ± SEM; n=4 mice per group Two-tailed unpaired Student's t-test, t=0.51, p=0.63. **H-M.** GFAP (**H**), IgG (**J**) and Claudin-5 (**L**) immunoreactivity in n-3 sufficient and n-3 deficient mice. Means ± SEM; n=3-5 mice per group. Two-tailed unpaired Student's t-test, t=0.18, p= 0.86, GFAP; t=1.364, p=0.215, IgG; t=1.718, p= 0.146, Claudin-5. **I,K,M.** Representative images of GFAP (**I**), IgG (**K**) and Claudin-5 (**M**) immunostainings. Arrows highlight claudin-5 staining.

**Supplementary Figure 4: Maternal n-3 PUFA deficiency modulates brain lipid composition.** Fatty acid composition of the cortex varies according to maternal n-3 PUFA intake. Means ± SEM; n=6 mice per group. Two-tailed unpaired Student’s t-test; t=11.94, ***p<0.0001, C18:1n-9; t=3.154, *p=0.01, C18:1n-7; t=7.27, ***p<0.0001, MUFAs; t=4.643, ***p=0.0009, AA; t=21.77, ***p<0.0001, C22:4n-6; t=36.09, ***p<0.0001, DPA n-6; t=35.78, ***p<0.0001, n-6 PUFAs; t=39.03, ***p<0.0001, DHA; t=39.36, ***p<0.0001, n-3 PUFAs; t=12.65, ***p<0.0001, C20:3n-9; t=5.371, ***p=0.0003, PUFAs; t=4.635, ***p=0.0009, Dimethylacetals (DMA) C16:0; t=4.271, **p=0.0016, DMA C18:1n-9; t=34.44, ***p<0.0001 n-6/n-3 ratio.

**Supplementary Figure 5: PS recognition may be involved in the elimination of spines in the developing hippocampus.** Pharmacological inhibition of the interaction between MFG-E8 and its microglial vitronectin receptor using a blocking peptide (cRGD) or its scrambled control (sc-cRGD) in standard diet-fed animals. Means ± SEM; n=4-5 mice per group. Two-tailed unpaired Student’s t-test, t=3.168, *p=0.016.

**Supplementary Figure 6: Maternal n-3 PUFA deficiency enhances the expression of complement protein C3 and of genes involved in microglia-mediated synaptic refinement.** *cd11b*, *cx3cr1*, *cx3cl1* and *tgfb* mRNA expression is increased in the hippocampus of n-3 deficient mice. Means ± SEM; n=5-9 mice per group. Two-tailed unpaired Student’s t-test; t=3.115, **p=0.0076, *cd11b*; t=4.58, **p=0.001, *cx3cr1*; t=3.916, **p=0.0016, *cx3cl1*; t=3.184, **p=0.0066*, tgfb*.

**Supplementary Figure 7:** **A.** Quantification of the percentage of microglial phagocytic cells 24h after application of AA. Means ± SEM; n=7-8 experiments per condition. Two-tailed unpaired Student’s t-test, Eth vs AA, t=0.543, p=0.596, LOW; t=0.134, p=0.89, MEDIUM; t=2.87, ^*^p=0.013, HIGH. **B.** Representative images of microglial cell in primary culture phagocyting latex beads. Scale bar=10µm. Arrows: highly phagocytic cells (>10 beads per cell body). **C.** Application of the CR3 antagonist XVA-143 significantly reduces the phagocytic activity of microglia towards beads. Means ± SEM; n=9 experiments per condition. Two-tailed unpaired Student’s t-test; t=2.216, *p=0.042. **D.** Representative images of AA/saline- and AA/XVA-treated microglial cells in primary culture. Scale bar=10µm. Arrowhead: highly phagocytic microglia.

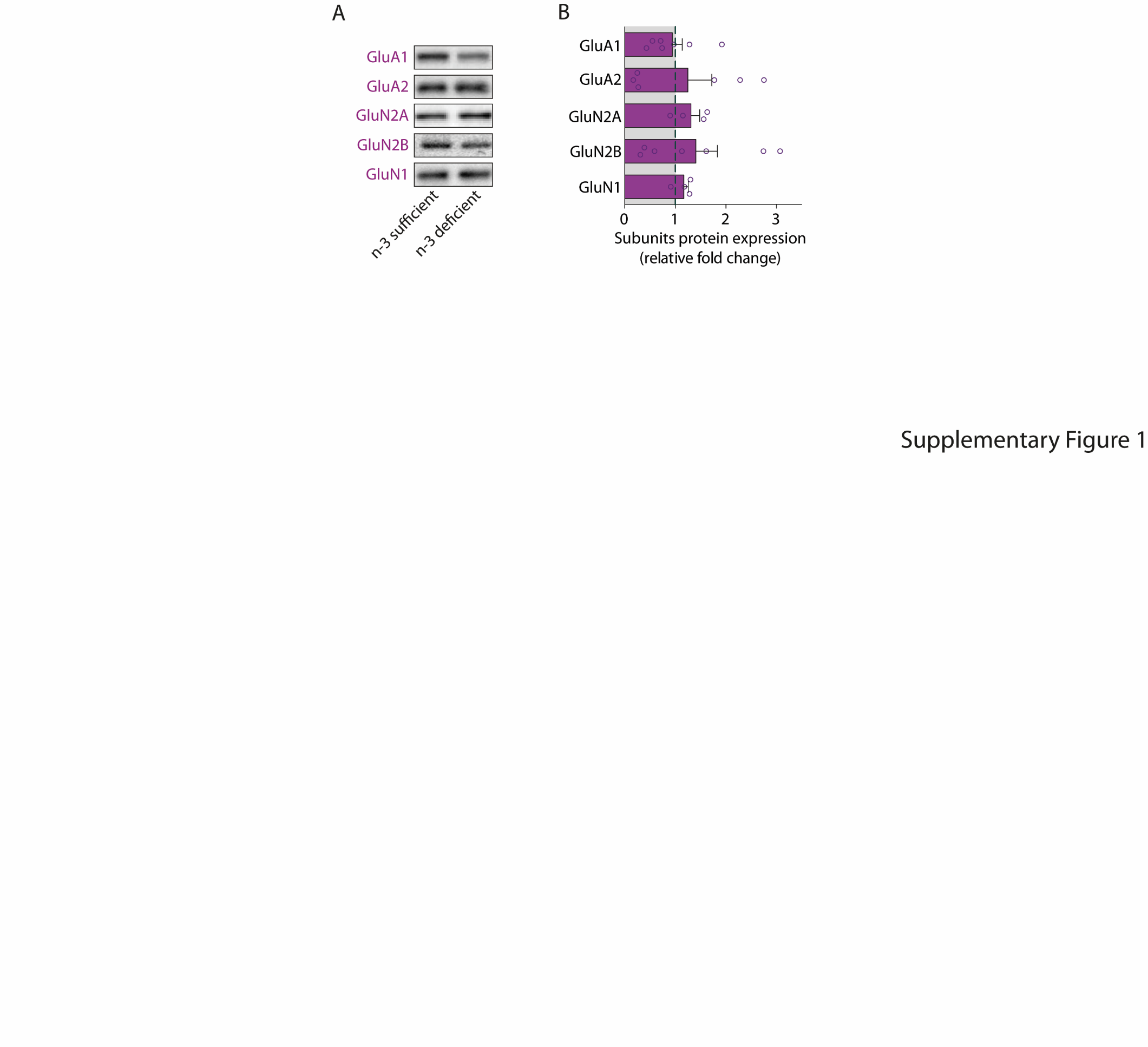

**Supplementary Figure 1**

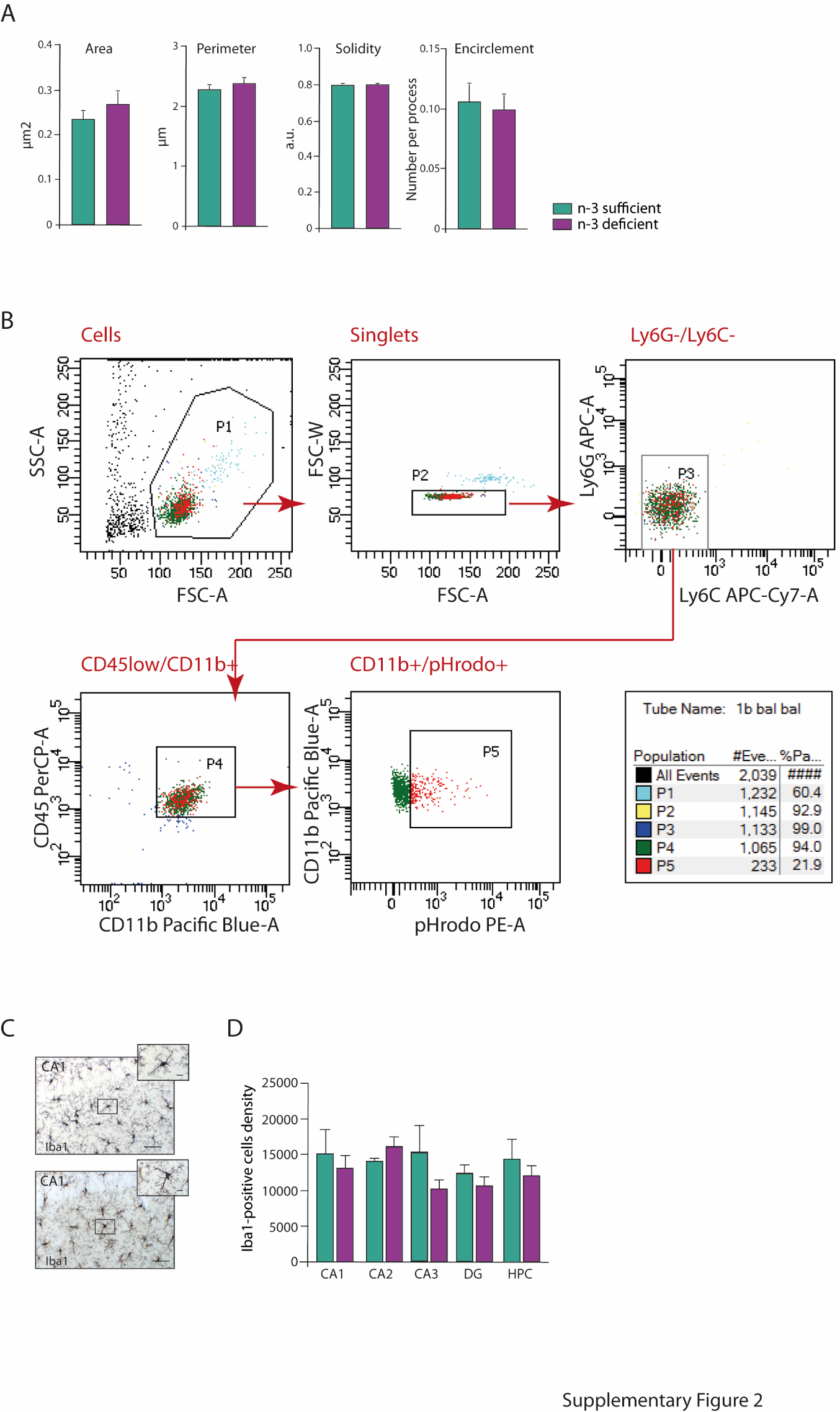

**Supplementary Figure 2**

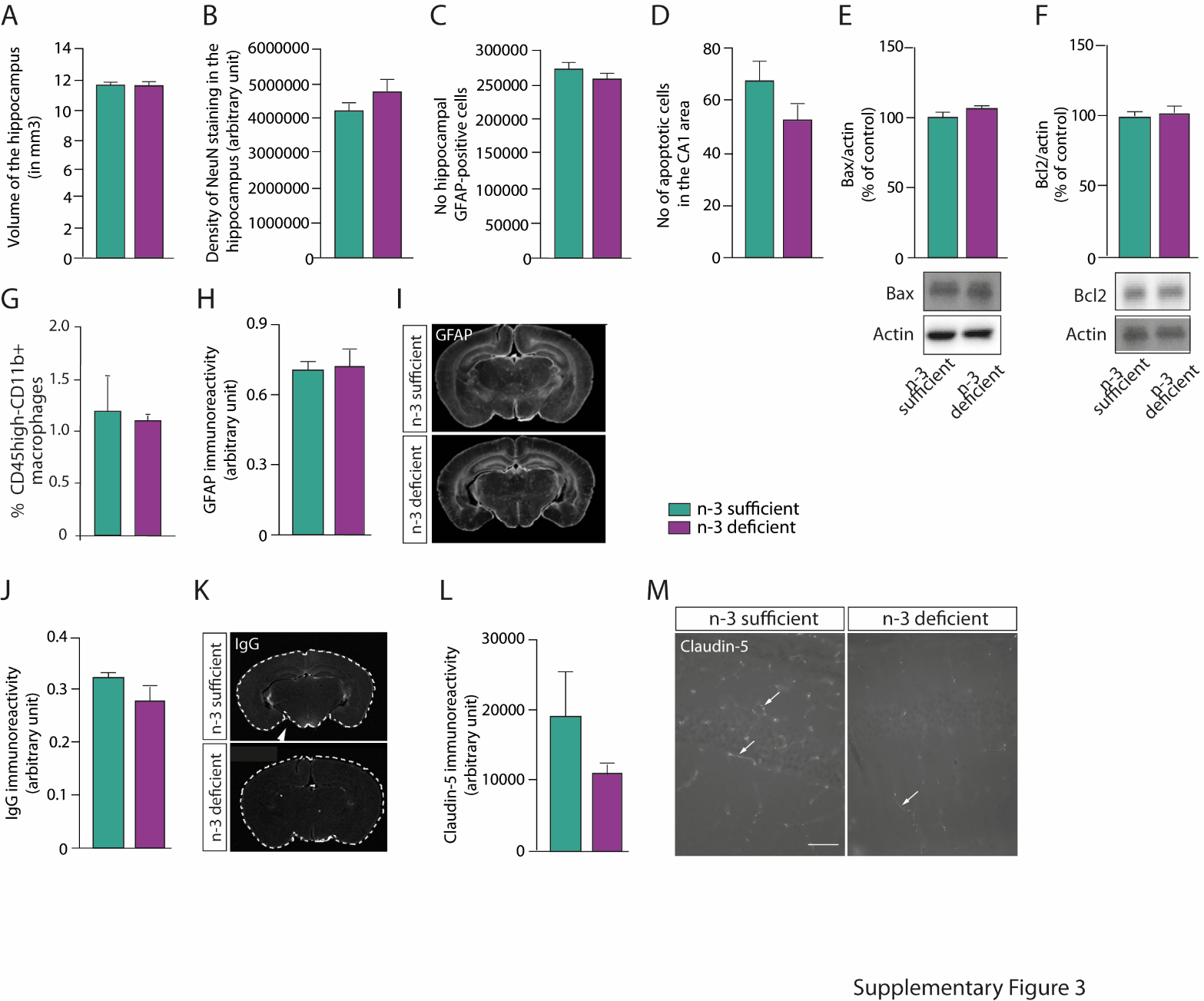

**Supplementary Figure 3**

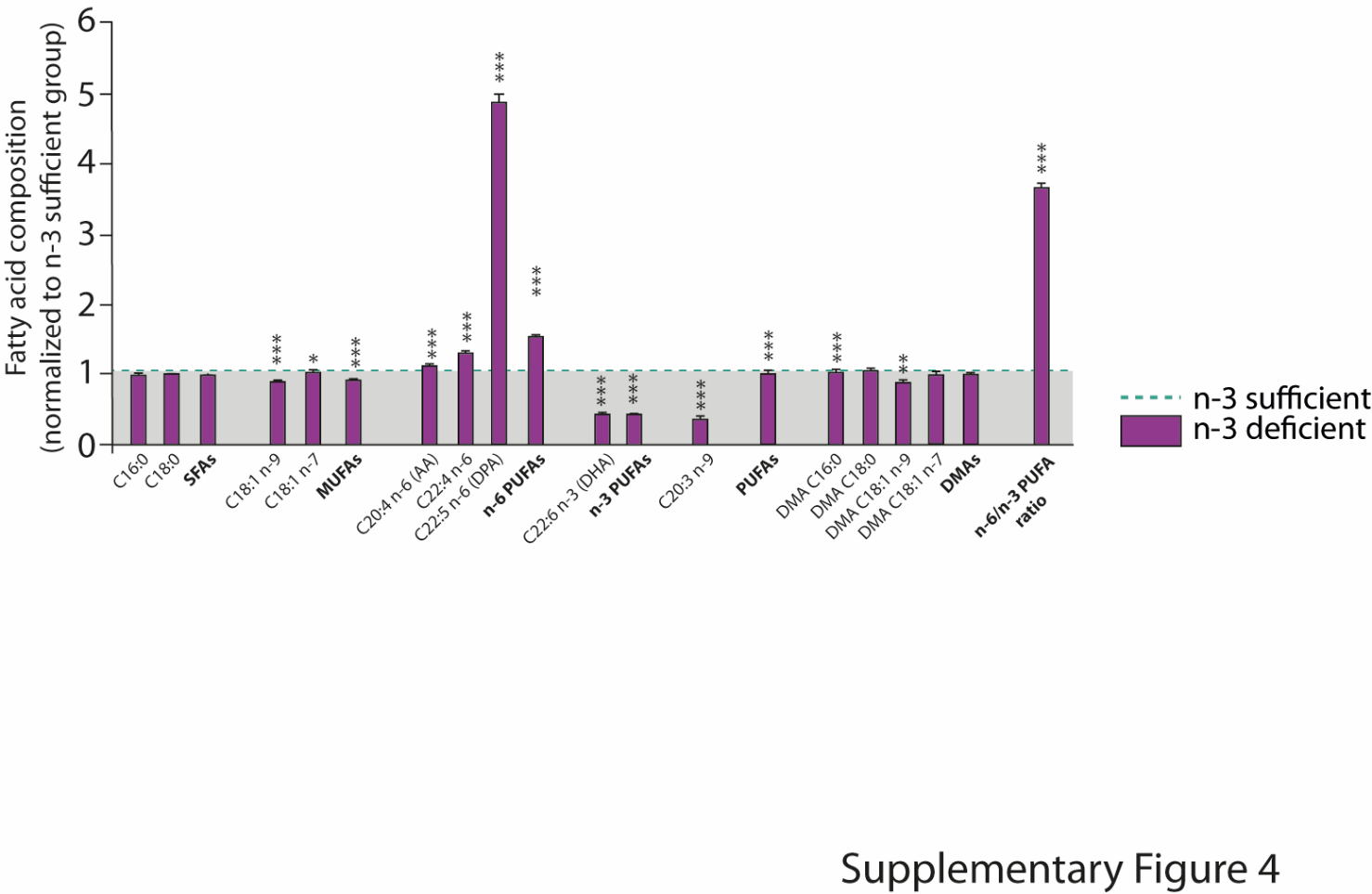

**Supplementary Figure 4**

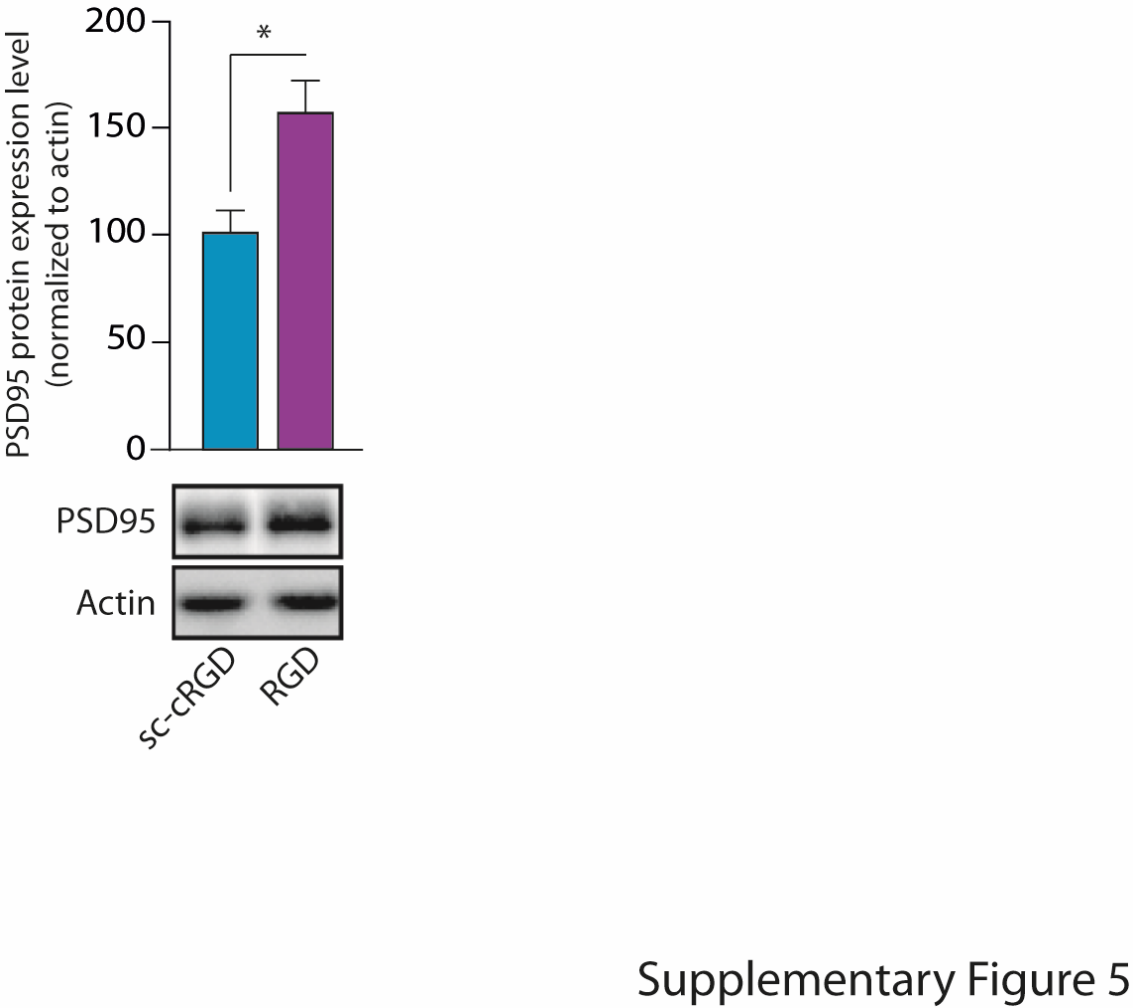

**Supplementary Figure 5**

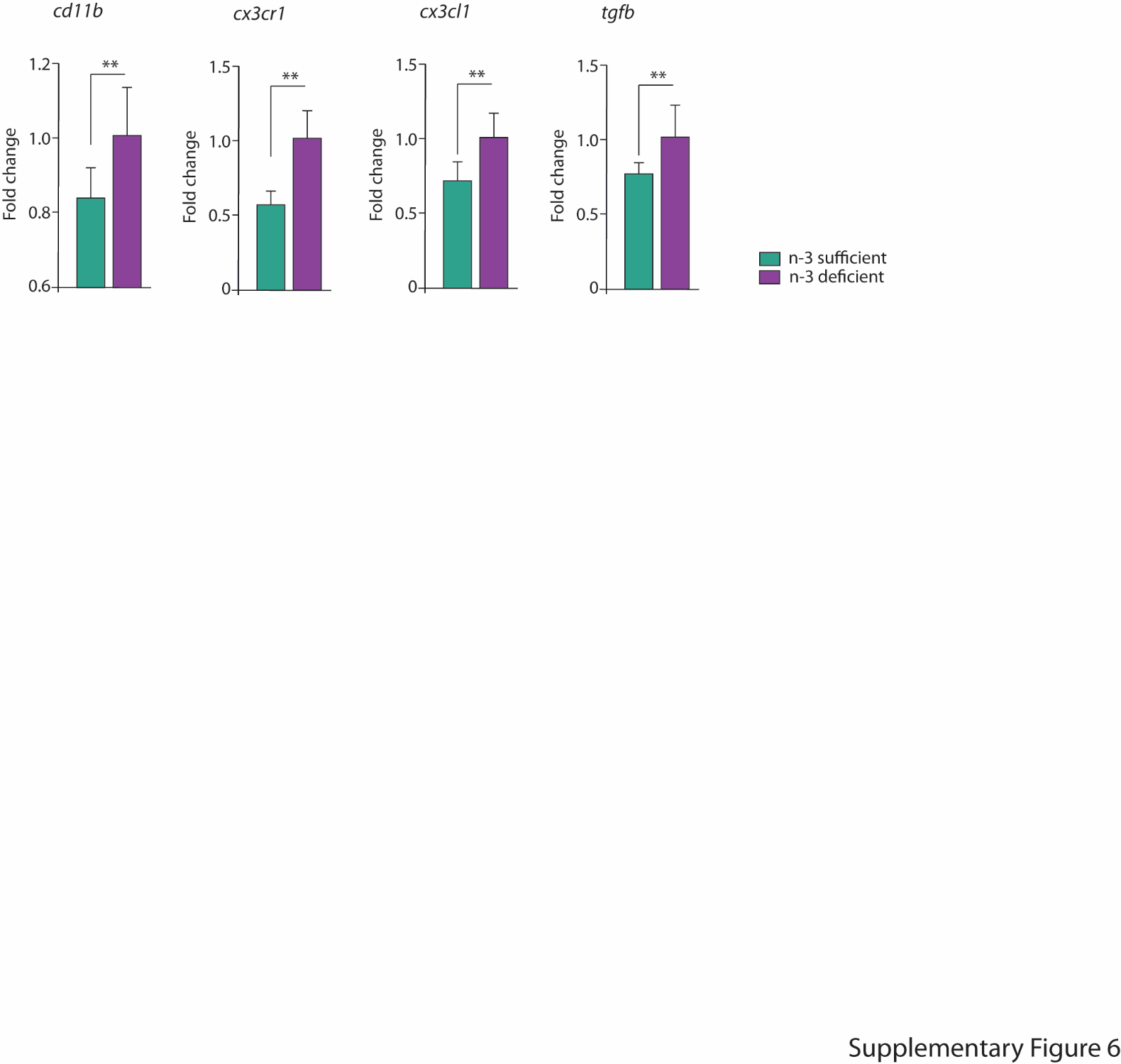

**Supplementary Figure 6**

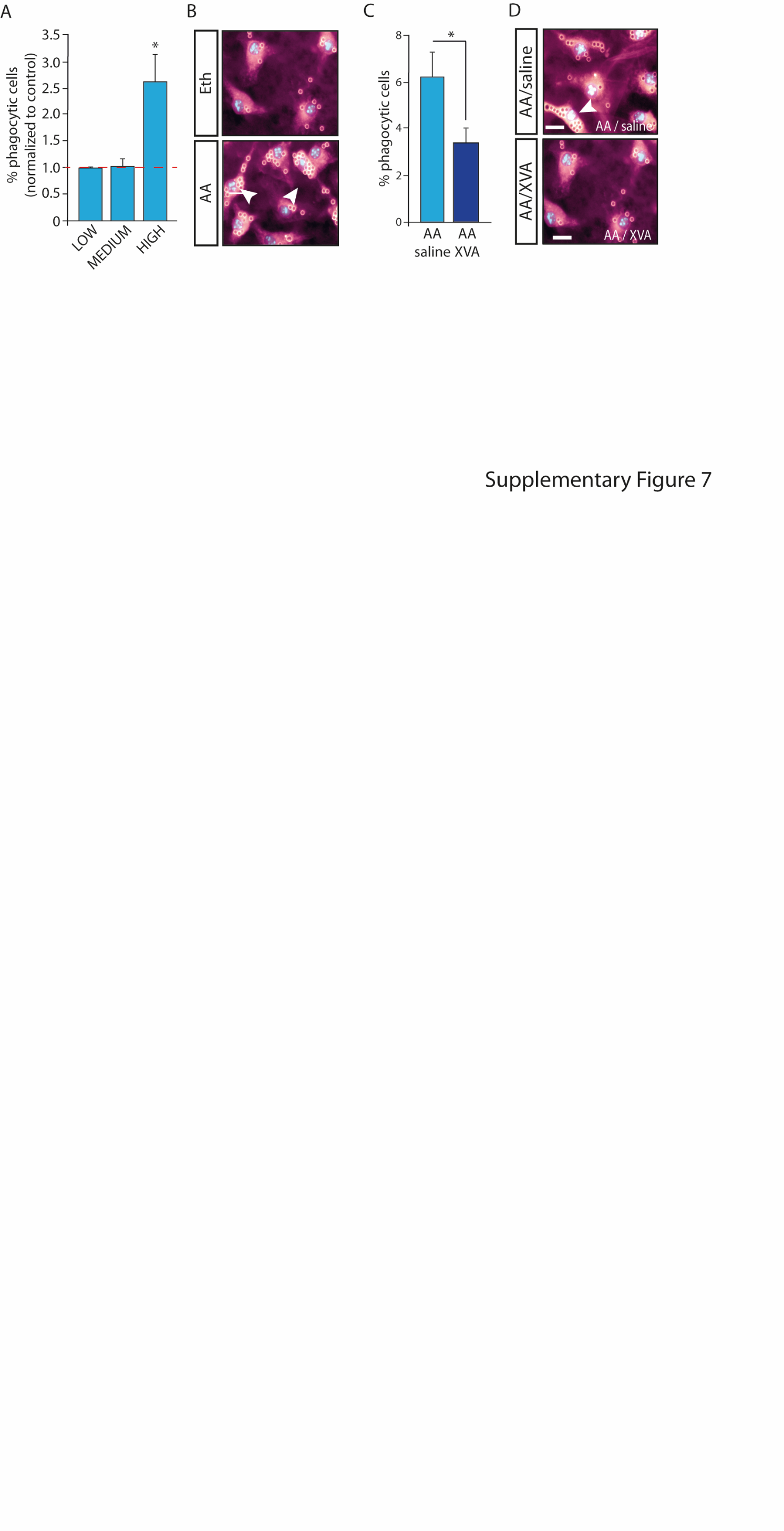

**Supplementary Figure 7**
