## Supplementary material for "Essential omega-3 fatty acids tune microglial phagocytosis of synaptic elements in the developing brain": Table 1

**Table 1: Composition of the diets (g/kg diet)**

| **Ingredient** | **Amount** |
| --- | --- |
| Casein | 180 |
| Cornstarch | 460 |
| Sucrose | 230 |
| Cellulose | 20 |
| Fat ^1^ | 50 |
| Mineral mix ^2^ | 50 |
| Vitamin mix ^3^ | 10 |

^1^: for detailed composition, see Table 2

^2^: composition (g/kg): sucrose, 110.7; CaCO_3_, 240; K_2_HPO_4_, 215; CaHPO_4_, 215; MgSO_4_,7H_2_O, 100; NaCl, 60; MgO, 40; FeSO_4_,7H_2_O, 8; ZnSO_4_,7H_2_O, 7; MnSO_4_,H_2_O, 2; CuSO_4_,5H_2_O, 1; Na_2_SiO_7_,3H_2_O, 0.5; AlK(SO_4_)_2_,12H_2_O, 0.2; K_2_CrO_4_, 0.15; NaF, 0.1; NiSO_4_,6H_2_O, 0.1; H_2_BO_3_, O.1; CoSO_4_,7H_2_O, 0.05; KIO_3_, 0.04; (NH_4_)_6_Mo_7_O_24_,4H_2_O, 0.02; LiCl, 0.015; Na_2_SeO_3_, 0.015; NH_4_VO_3_, 0.01

^3^: composition (g/kg): sucrose, 549.45; retinyl acetate, 1; cholecalciferol, 0.25; DL-α-tocopheryl acetate, 20; phylloquinone, 0.1; thiamin HCl, 1; riboflavin, 1; nicotinic acid, 5; calcium pantothenate, 2.5; pyridoxine HCl, 1; biotin, 1; folic acid, 0.2; cyanobalamin, 2.5; choline HCl, 200; DL-methionin, 200; p-aminobenzoic acid, 5; inositol, 10
