## Supplementary material for "Essential omega-3 fatty acids tune microglial phagocytosis of synaptic elements in the developing brain": Table 2

**Table 2: Fatty acid composition of the diets (% wt of total fatty acids)**

| diets | Deficient | Sufficient | Standard chow  (A04) |
| --- | --- | --- | --- |
| 16:0 | 7.3 | 22.6 | 20.3 |
| 18:0 | 4.1 | 3.3 | 2.2 |
| other saturated FAs | 1.6 | 1.8 | 1.8 |
| total saturated FAs | 13.0 | 27.7 | 24.3 |
| 18:1n-9 | 28.1 | 57.9 | 19.3 |
| 18:1n-7 | 0.9 | 1.5 | 1.5 |
| other monounsaturated FAs | 0.2 | 0.4 | 3.9 |
| total monounsaturated FAs | 29.4 | 60.0 | 24.7 |
| 18:2n-6 (LA) | 57.4 | 10.6 | 45.9 |
| Other n-6 PUFAs | n.d. | n.d. | 0.3 |
| total n-6 PUFAs | 57.4 | 10.7 | 46.2 |
| 18:3n-3 (ALA) | 0.2 | 1.6 | 3.3 |
| 20:5 n-3 | n.d. | n.d. | 0.6 |
| 22:5 n-3 | n.d. | n.d. | 0.1 |
| 22:6 n-3 | n.d. | n.d. | 0.8 |
| total n-3 PUFAs | 0.2 | 1.6 | 4.8 |
| total PUFAs | 57.6 | 12.3 | 51.0 |
| LA/ALA | 287 | 6.6 | 13.9 |

FAs, fatty acids; PUFAs, polyunsaturated fatty acids; LA: linoleic acid; ALA, α-linolenic acid; n.d., not detected (under the limit for the detection by gas chromatography, <0.05%).
