## Supplementary material for "Essential omega-3 fatty acids tune microglial phagocytosis of synaptic elements in the developing brain": Table 3

| **Gene name** | **Ratio**  **n-3 deficient/**  **n-3 sufficient** |  |  |  |  |
| --- | --- | --- | --- | --- | --- |
| **Ccl3** | 50,2 | **Slco2b1** | 0,55 | **Ttr** | 0,35 |
| **Ccl4** | 18,9 | **Golm1** | 0,55 | **BMP6** | 0,35 |
| **Rasgef1b** | 9,5 | **A430107D22Rik** | 0,55 | **NOX1** | 0,34 |
| **Cd83** | 5,4 | **D830030K20Rik** | 0,54 | **Nav2** | 0,34 |
| **Atf3** | 5,3 | **Vat1l** | 0,54 | **Mmp12** | 0,34 |
| **Ccrl2** | 4,7 | **LOC100038847** | 0,54 | **Hist1h2ac** | 0,34 |
| **C3ar1** | 4,4 | **Ifi202b** | 0,54 | **Siglec1** | 0,33 |
| **Ccl2** | 4,1 | **Rgmb** | 0,54 | **Plxna1** | 0,32 |
| **Ppp1r15a** | 3,5 | **Ccr5** | 0,54 | **Fcrls** | 0,32 |
| **Nlrp3** | 3,0 | **ADAM17** | 0,54 | **Slco4a1** | 0,31 |
| **Apoe** | 3,0 | **Ppp1r9a** | 0,53 | **Cbr3** | 0,31 |
| **Rgs1** | 2,9 | **Rnf180** | 0,53 | **Slc46a1** | 0,31 |
| **Trim47** | 2,8 | **FcgrIIIa (CD16a)** | 0,53 | **Upk1b** | 0,29 |
| **Adamts1** | 2,8 | **P2ry12** | 0,53 | **Khdrbs3** | 0,29 |
| **Dab2** | 2,7 | **CD200R** | 0,52 | **Tnfrsf17** | 0,29 |
| **Klhl38** | 2,6 | **Gpr165** | 0,52 | **TREM1** | 0,28 |
| **1810011O10Rik** | 2,6 | **Fgd2** | 0,52 | **ll7r** | 0,28 |
| **CEBPB** | 2,5 | **Trem2** | 0,51 | **Ak1** | 0,28 |
| **Myc** | 2,4 | **P4ha1** | 0,51 | **BMP7** | 0,28 |
| **Cd14** | 2,4 | **Gtf2h2** | 0,51 | **WNT5A** | 0,27 |
| **Fos** | 2,3 | **Zfp691** | 0,51 | **Kitl** | 0,24 |
| **Il1a** | 2,3 | **Tmem144** | 0,51 | **Chi3l1** | 0,22 |
| **Rgl1** | 2,2 | **Plxna4** | 0,51 | **Cfb** | 0,21 |
| **Fosb** | 2,2 | **Itgax** | 0,51 | **Arg1** | 0,20 |
| **Manba** | 2,2 | **Bend6** | 0,51 | **Olfml3** | 0,18 |
| **Npl** | 2,1 | **Csf3r** | 0,51 | **Gal3st4** | 0,17 |
| **cathepsin** | 2,0 | **SEMA3C** | 0,50 |  |  |
| **Rhob** | 2,0 | **Bco2** | 0,50 |  |  |
| **Ltc4s** | 2,0 | **MPO (myeloperoxidase)** | 0,50 |  |  |
| **Egr1** | 2,0 | **ENSMUSG00000079376** | 0,50 |  |  |
| **Fth1** | 1,9 | **Erf** | 0,50 |  |  |
| **Abca1** | 1,9 | **Tmc7** | 0,49 |  |  |
| **Jun** | 1,9 | **4933406P04Rik** | 0,49 |  |  |
| **Socs3** | 1,8 | **Mmp2** | 0,48 |  |  |
| **Junb** | 1,7 | **Tmem204** | 0,48 |  |  |
| **SREBP1** | 1,7 | **Tlr3** | 0,48 |  |  |
| **Tlr2** | 1,6 | **Spnb4** | 0,47 |  |  |
| **Il1rl2** | 1,5 | **Scoc** | 0,47 |  |  |
| **Ctsl** | 1,5 | **B930046C15Rik** | 0,47 |  |  |
| **9030625A04Rik** | 1,4 | **Asph** | 0,47 |  |  |
| **SRA1** | 1,4 | **WNT7A** | 0,47 |  |  |
|  |  | **Pycard** | 0,46 |  |  |
| **Icam1** | 0,82 | **Sall1** | 0,46 |  |  |
| **HIST1H2AB** | 0,76 | **Pros1** | 0,46 |  |  |
| **Ryk** | 0,75 | **Garnl3** | 0,46 |  |  |
| **Grm1** | 0,75 | **Fabp5** | 0,45 |  |  |
| **Rtn1** | 0,75 | **MR** | 0,45 |  |  |
| **Tfeb** | 0,74 | **Camk2n1** | 0,44 |  |  |
| **Map3k7** | 0,72 | **B4galt4** | 0,44 |  |  |
| **Slc24a3** | 0,71 | **Slc2a5** | 0,44 |  |  |
| **Qdpr** | 0,70 | **Il21r** | 0,43 |  |  |
| **ADAM10** | 0,68 | **IL34** | 0,43 |  |  |
| **Cxxc5** | 0,67 | **Atp8a2** | 0,43 |  |  |
| **Ckb** | 0,67 | **Jam2** | 0,43 |  |  |
| **Ttc28** | 0,66 | **Myo1b** | 0,42 |  |  |
| **Tmem100** | 0,66 | **Gpr56** | 0,42 |  |  |
| **Nfia** | 0,65 | **Ebf3** | 0,42 |  |  |
| **Tm9sf4** | 0,65 | **Olfml2b** | 0,42 |  |  |
| **Snn** | 0,64 | **Fgfr1** | 0,42 |  |  |
| **Abi3** | 0,64 | **Tmem119** | 0,42 |  |  |
| **Epn2** | 0,64 | **Scamp5** | 0,42 |  |  |
| **Sema4d** | 0,63 | **Fads1** | 0,41 |  |  |
| **Map2k1** | 0,62 | **Psd** | 0,41 |  |  |
| **Zfpm1** | 0,62 | **Tmeff1** | 0,41 |  |  |
| **Rbbp9** | 0,62 | **Ptprm** | 0,41 |  |  |
| **Adora3** | 0,62 | **Tlr5** | 0,40 |  |  |
| **Ang** | 0,61 | **Adamts16** | 0,40 |  |  |
| **Gpr34** | 0,61 | **Eng** | 0,40 |  |  |
| **Rtn4rl1** | 0,61 | **CD36** | 0,39 |  |  |
| **Bin1** | 0,60 | **Itga9** | 0,39 |  |  |
| **Acp2** | 0,60 | **Gpr40** | 0,39 |  |  |
| **Epb4.1l2** | 0,60 | **Tspan18** | 0,39 |  |  |
| **Cttnbp2nl** | 0,60 | **Ecscr** | 0,39 |  |  |
| **Itgb5** | 0,59 | **Cntn1** | 0,39 |  |  |
| **Sgce** | 0,58 | **WNT2** | 0,39 |  |  |
| **D18Ertd653e** | 0,58 | **Gp9** | 0,38 |  |  |
| **Gm10790** | 0,57 | **Arhgap22** | 0,37 |  |  |
| **Tjp1** | 0,57 | **Pon3** | 0,37 |  |  |
| **Spsb1** | 0,57 | **Plxdc2** | 0,37 |  |  |
| **C3** | 0,57 | **Tanc2** | 0,37 |  |  |
| **Lair1** | 0,57 | **Rab6b** | 0,36 |  |  |
| **Tppp** | 0,57 | **Kcnd1** | 0,36 |  |  |
| **IL6ST** | 0,56 | **Csmd3** | 0,36 |  |  |
| **Cx3cr1** | 0,56 | **NRG1** | 0,36 |  |  |
| **Siglech** | 0,56 | **Capn3** | 0,35 |  |  |

**Table 3 : List of all genes that are significantly modulated by maternal n-3 PUFA deficiency.** Only genes with P<0.05 are considered significant. In orange: down-regulated genes, in blue: upregulated genes.
